# Eukaryotic virus composition can predict the efficiency of carbon export in the global ocean

**DOI:** 10.1101/710228

**Authors:** Hiroto Kaneko, Romain Blanc-Mathieu, Hisashi Endo, Samuel Chaffron, Tom O. Delmont, Morgan Gaia, Nicolas Henry, Rodrigo Hernández-Velázquez, Canh Hao Nguyen, Hiroshi Mamitsuka, Patrick Forterre, Olivier Jaillon, Colomban de Vargas, Matthew B. Sullivan, Curtis A. Suttle, Lionel Guidi, Hiroyuki Ogata

## Abstract

The biological carbon pump, in which carbon fixed by photosynthesis is exported to the deep ocean through sinking, is a major process in Earth’s carbon cycle. The proportion of primary production that is exported is termed the carbon export efficiency (CEE). Based on in-lab or regional scale observations, viruses were previously suggested to affect the CEE (i.e., viral “shunt” and “shuttle”). In this study, we tested associations between viral community composition and CEE measured at a global scale. A regression model based on relative abundance of viral marker genes explained 67% of the variation in CEE. Viruses with high importance in the model were predicted to infect ecologically important hosts. These results are consistent with the view that the viral shunt and shuttle functions at a large scale and further imply that viruses likely act in this process in a way dependent on their hosts and ecosystem dynamics.

## Introduction

A major process in the global cycling of carbon is the oceanic biological carbon pump (BCP), an organism-driven process by which atmospheric carbon (*i.e.,* CO_2_) is transferred and sequestered to the ocean interior and seafloor for periods ranging from centuries to hundreds of millions of years. Between 15% and 20% of net primary production (NPP) is exported out of the euphotic zone, with 0.3% of fixed carbon reaching the seafloor annually (Zhang et al., 2018). However, there is wide variation in estimates of the proportion of primary production in the surface ocean that is exported to depth, ranging from 1% in the tropical Pacific to 35-45% during the North Atlantic bloom (Buesseler and Boyd, 2009). As outlined below, many factors affect the BCP.

Of planktonic organisms living in the upper layer of the ocean, diatoms (Tréguer et al., 2018) and zooplankton (Turner, 2015) have been identified as important contributors to the BCP in nutrient-replete oceanic regions. In the oligotrophic ocean, cyanobacteria, collodarians (Lomas and Moran, 2011), diatoms (Agusti et al., 2015; Karl et al., 2012; Leblanc et al., 2018), and other small (pico-to nano-) plankton (Lomas and Moran, 2011) have been implicated in the BCP. Sediment trap studies suggest that ballasted aggregates of plankton with biogenic minerals contribute to carbon export to the deep sea (Iversen and Ploug, 2010; Klaas and Archer, 2002). The BCP comprises three processes: carbon fixation, export, and remineralization. As these processes are governed by complex interactions between numerous members of planktonic communities (Zhang et al., 2018), the BCP is expected to involve various organisms, including viruses (Zimmerman et al., 2019a).

Viruses have been suggested to regulate the efficiency of the BCP. Lysis of host cells by viruses releases cellular material in the form of dissolved organic matter (DOM), which fuels the microbial loop and enhances respiration and secondary production (Gobler et al., 1997; Weitz et al., 2015). This process, coined “viral shunt (Wilhelm and Suttle, 1999)”, can reduce the carbon export efficiency (CEE) because it increases the retention of nutrients and carbon in the euphotic zone and prevents their transfer to higher trophic levels as well as their export from the euphotic zone to the deep sea (Fuhrman, 1999; Weitz et al., 2015). However, an alternative process is also considered, in which viruses contribute to the vertical carbon export (Weinbauer, 2004). For instance, a theoretical study proposed that the CEE increases if viral lysis augments the ratio of exported carbon relative to the primary production-limiting nutrients (nitrogen and phosphorous) (Suttle, 2007). Laboratory experimental studies reported that cells infected with viruses form larger particles (Peduzzi and Weinbauer, 1993; Yamada et al., 2018), can sink faster (Lawrence and Suttle, 2004), and can lead to preferential grazing by heterotrophic protists (Evans and Wilson, 2008) and/or to higher growth of grazers (Goode et al., 2019). This process termed “viral shuttle (Sullivan et al., 2017)” is supported by several field studies that reported association of viruses with sinking material. Viruses were observed in sinking material in the North Atlantic Ocean (Proctor and Fuhrman, 1991) and sediment of coastal waters where algal blooms occur (Lawrence et al., 2002; Tomaru et al., 2007, 2011). In addition, vertical transport of bacterial viruses between photic and aphotic zones was observed in the Pacific Ocean (Hurwitz et al., 2015) and in *Tara* Oceans virome data (Brum et al., 2015). A systematic analysis of large-scale omics data from oligotrophic oceanic regions revealed a strong positive association between carbon flux and bacterial dsDNA viruses (*i.e.*, cyanophages), which were previously unrecognized as possible contributors to the BCP (Guidi et al., 2016). More recently, viral infection of blooms of the photosynthetic eukaryote *Emiliania huxleyi* in the North Atlantic were found to be accompanied by particle aggregation and greater downward vertical flux of carbon, with the highest export during the early stage of viral infection (Laber et al., 2018). These studies raise the question of the overall impact of viruses infecting eukaryotes on oceanic carbon cycling and export. Given the significant contributions of eukaryotic plankton to ocean biomass and net production (Hirata et al., 2011; Li, 1995) and their observed predominance over prokaryotes in sinking materials of Sargasso Sea oligotrophic surface waters (Fawcett et al., 2011; Lomas and Moran, 2011), various lineages of eukaryotic viruses may be responsible for a substantial part of the variation in carbon export across oceanic regions.

If the “viral shunt” and “shuttle” processes function at a global scale and if these involve specific viruses, we expect to detect a statistical association between viral community composition and CEE in a large scale omics data. To our knowledge, such an association has never been investigated. Although this test per se does not prove that viruses regulate CEE, we consider the association is worth being tested because such an association is a necessary condition for the global model of viral shunt and shuttle and, under its absence, we will have to reconsider the model. Deep sequencing of planktonic community DNA and RNA, as carried out in *Tara* Oceans, has enabled the identification of marker genes of major viral groups infecting eukaryotes (Hingamp et al., 2013; Carradec et al., 2018; Culley, 2018). To examine the association between viral community composition and CEE, we thus used the comprehensive organismal dataset from the *Tara* Oceans expedition (Carradec et al., 2018; Sunagawa et al., 2015), as well as related measurements of carbon export estimated from particle concentrations and size distributions observed *in situ* (Guidi et al., 2016).

In the present study, we identified several hundred marker-gene sequences of nucleocytoplasmic large DNA viruses (NCLDVs) in metagenomes of 0.2–3 μm size fraction. We also identified RNA and ssDNA viruses in metatranscriptomes of four eukaryotic size fractions spanning 0.8 to 2,000 μm. The resulting profiles of viral distributions were compared with an image-based measure of carbon export efficiency (CEE), which is defined as the ratio of the carbon flux at depth to the carbon flux at surface.

## Results and Discussion

### Detection of diverse eukaryotic viruses in *Tara* Oceans gene catalogs

We used profile hidden Markov model-based homology searches to identify marker-gene sequences of eukaryotic viruses in two ocean gene catalogs. These catalogs were previously constructed from environmental shotgun sequence data of samples collected during the *Tara* Oceans expedition. The first catalog, the Ocean Microbial Reference Gene Catalog (OM-RGC), contains 40 million non-redundant genes predicted from the assemblies of *Tara* Oceans viral and microbial metagenomes (Sunagawa et al., 2015). We searched this catalog for NCLDV DNA polymerase family B (PolB) genes, as dsDNA viruses may be present in microbial metagenomes because large virions (> 0.2 μm) have been retained on the filter or because viral genomes actively replicating or latent within picoeukaryotic cells have been captured. The second gene catalog, the Marine Atlas of *Tara* Oceans Unigenes (MATOU), contains 116 million non-redundant genes derived from metatranscriptomes of single-cell microeukaryotes and small multicellular zooplankton (Carradec et al., 2018). We searched this catalog for NCLDV PolB genes, RNA-dependent RNA polymerase (RdRP) genes of RNA viruses, and replication-associated protein (Rep) genes of ssDNA viruses, since transcripts of viruses actively infecting their hosts, as well as genomes of RNA viruses, have been captured in this catalog.

We identified 3,874 NCLDV PolB sequences (3,486 in metagenomes and 388 in metatranscriptomes), 975 RNA virus RdRP sequences, and 299 ssDNA virus Rep sequences (Table 1). These sequences correspond to operational taxonomic units (OTUs) at a 95% identity threshold. All except 17 of the NCLDV PolBs from metagenomes were assigned to the families *Mimiviridae* (*n* = 2,923), *Phycodnaviridae* (*n* = 348), and *Iridoviridae* (*n* = 198) (Table 1). The larger numbers of PolB sequences assigned to *Mimiviridae* and *Phycodnaviridae* compared with other NCLDV families are consistent with a previous observation based on a smaller dataset (Hingamp et al., 2013). The divergence between these environmental sequences and reference sequences from known viral genomes was greater in *Mimiviridae* than in *Phycodnaviridae* (Figure 1a, S1a and S2). Within *Mimiviridae,* 83% of the sequences were most similar to those from algae-infecting *Mimivirus* relatives. Among the sequences classified in *Phycodnaviridae,* 93% were most similar to those in *Prasinovirus*, whereas 6% were closest to *Yellowstone lake phycodnavirus*, which is closely related to *Prasinovirus*. Prasinoviruses are possibly over-represented in the metagenomes because the 0.2 to 3 μm size fraction selects their picoeukaryotic hosts. RdRP sequences were assigned mostly to the order *Picornavirales* (*n* = 325), followed by the families *Partitiviridae* (*n* = 131), *Narnaviridae* (*n* = 95), *Tombusviridae* (*n* = 45), and *Virgaviridae* (*n* = 33) (Table 1), with most sequences being distant (30% to 40% amino acid identity) from reference viruses (Figures 1b, S1b and S3). These results are consistent with previous studies on the diversity of marine RNA viruses, in which RNA virus sequences were found to correspond to diverse positive-polarity ssRNA and dsRNA viruses distantly related to well-characterized viruses (Culley, 2018). *Picornavirales* may be over-represented in the metatranscriptomes because of the polyadenylated RNA selection. The majority (*n* = 201) of Rep sequences were annotated as *Circoviridae*, known to infect animals, which is consistent with a previous report (Wang et al., 2018). Only eight were annotated as plant ssDNA viruses (families *Nanovoridae* and *Gemniviridae*) (Table 1). Most of these environmental sequences are distant (40% to 50% amino acid identity) from reference sequences (Figures 1c, S1c and S4). Additional 388 NCLDV PolBs were detected in the metranscriptomes. The average cosmopolitanism (number of samples where an OTU was observed by at least two reads) for PolBs in metagenomes was 23 samples against 2.9 for metranscriptome-derived PolB sequences, 5.5 for Reps, and 5.8 for RdRPs. Within metatranscriptomes, the average gene-length normalized read counts for PolBs were respectively ten and three times lower than those of RdRPs and Reps. Therefore, PolBs from metatranscriptomes were not further used in our study.

**Figure 1:**
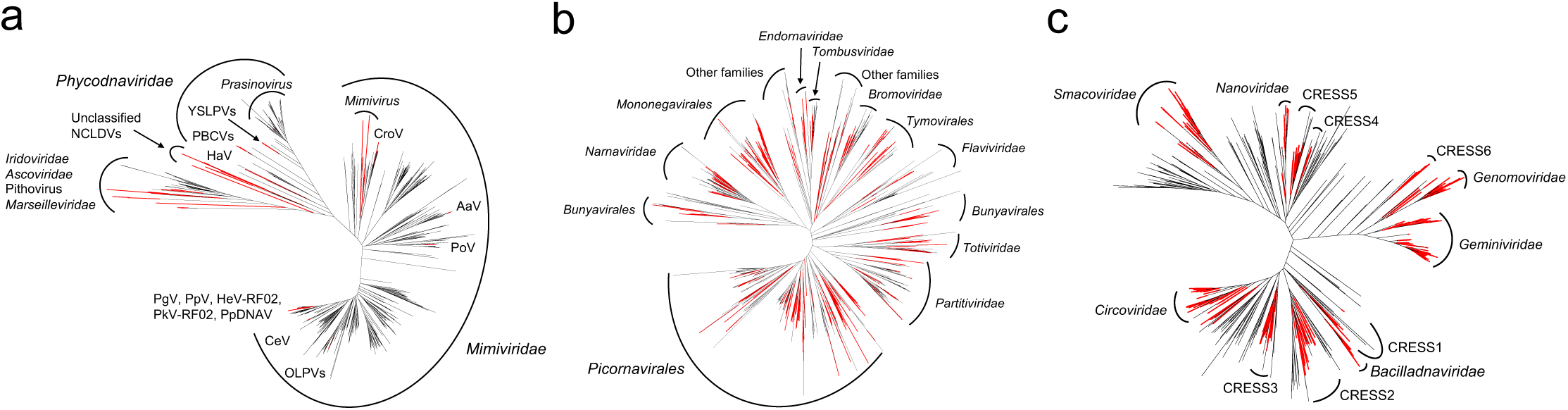
Viruses of eukaryotic plankton identified in *Tara* Oceans samples are distantly related to characterized viruses. Unrooted maximum likelihood phylogenetic trees containing environmental (black) and reference (red) viral sequences for NCLDV DNA polymerase family B (**a**), RNA virus RNA-dependent RNA polymerase (**b**), and ssDNA virus replication-associated protein (**c**). A rectangular representation of these trees with branch support values is provided in Figure S2–S4.

**Table 1:**
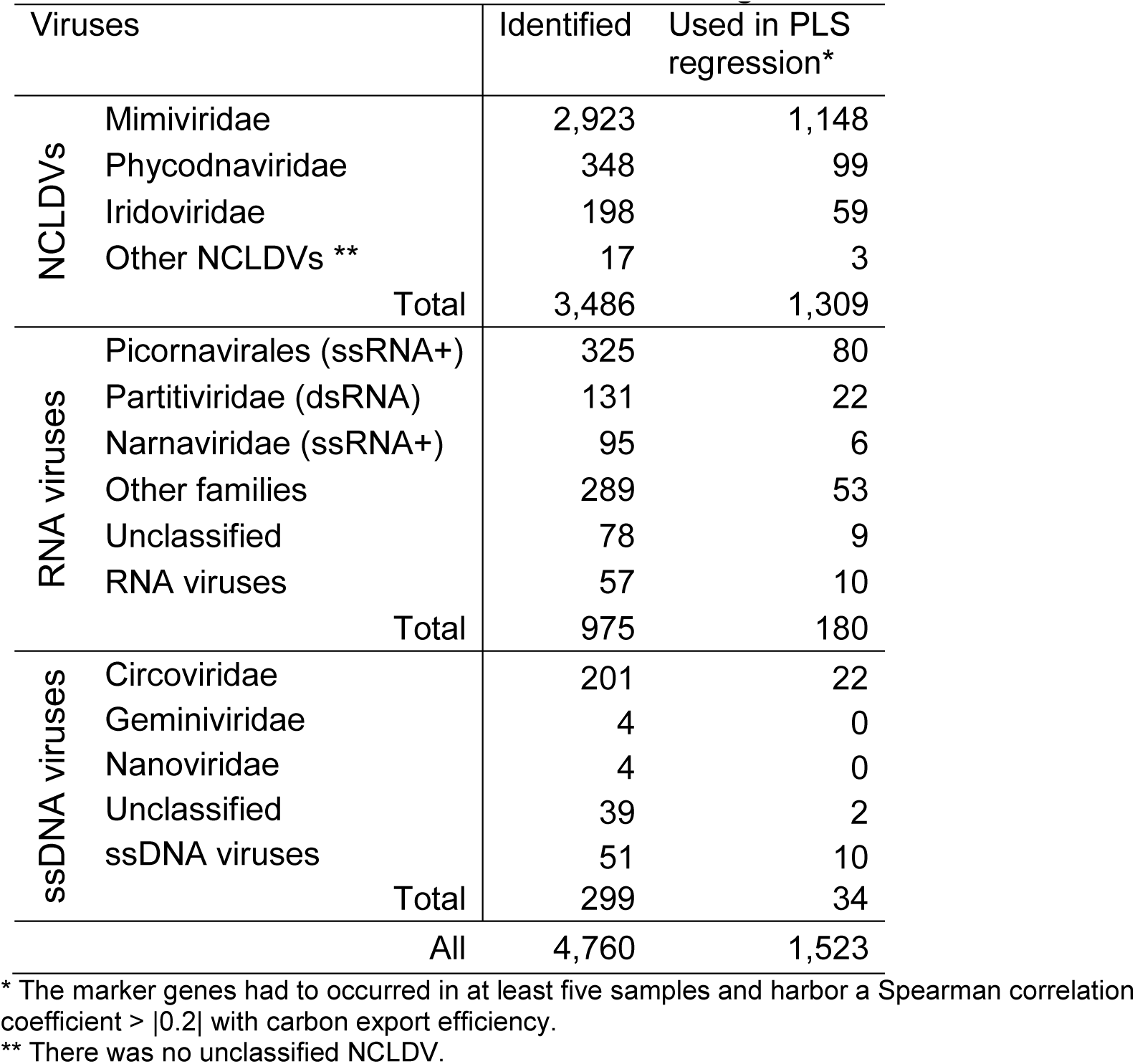
Taxonomic breakdown of viral marker genes

### Composition of eukaryotic viruses can explain the variation of carbon export efficiency

Among the PolB, RdRP, and Rep sequences identified in the *Tara* Oceans gene catalogs, 38%, 18%, and 11% (total = 1,523 sequences), respectively, were present in at least five samples and had matching carbon export measurement data (Table 1). We used the relative abundance (defined as the centered log-ratio transformed gene-length normalized read count) profiles of these 1,523 marker-gene sequences at 59 sampling sites in the photic zone of 39 *Tara* Oceans stations (Figure 2) to test for association between their composition and a measure of carbon export efficiency (CEE, see Transparent Methods, Figure S5). A partial least squares (PLS) regression model explained 67% (coefficient of determination *R*^2^ = 67%) of the variation in CEE with a Pearson correlation coefficient of 0.84 between observed and predicted values. This correlation was confirmed to be statistically significant by permutation test (*P* < 1 × 10^−4^) (Figure 3a).

**Figure 2:**
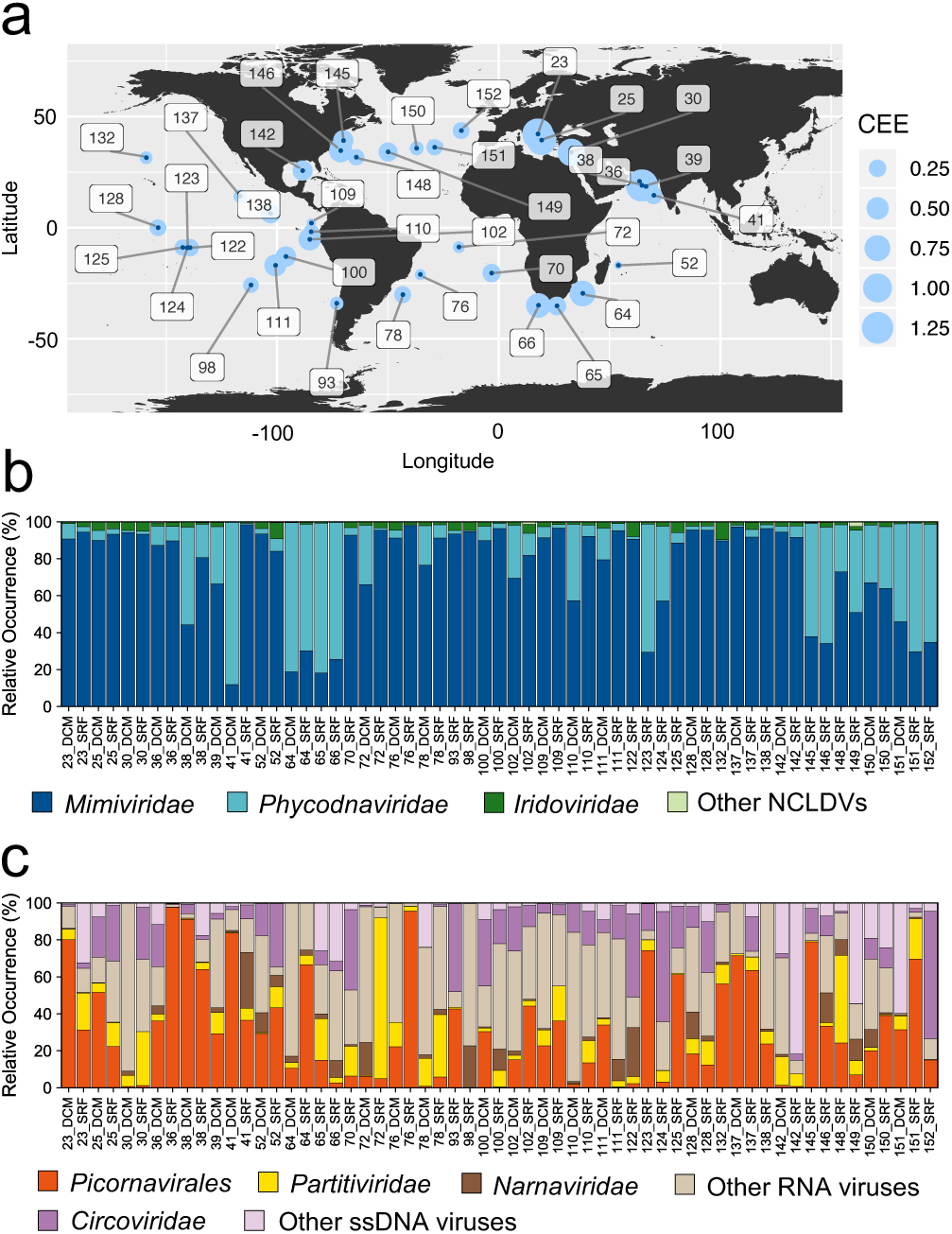
Carbon export efficiency and relative marker-gene occurrence of eukaryotic plankton viruses along the sampling route. **a** Carbon export efficiency estimated at 39 *Tara* Oceans stations where surface and DCM layers were sampled for prokaryote-enriched metagenomes and eukaryotic metatranscriptomes. **b and c** Relative marker-gene occurrence of major groups of viruses of eukaryotic plankton for NCLDVs in metagenomes (**b**) and for RNA and ssDNA viruses in metatranscriptomes (**c**) at 59 sampling sites.

**Figure 3:**
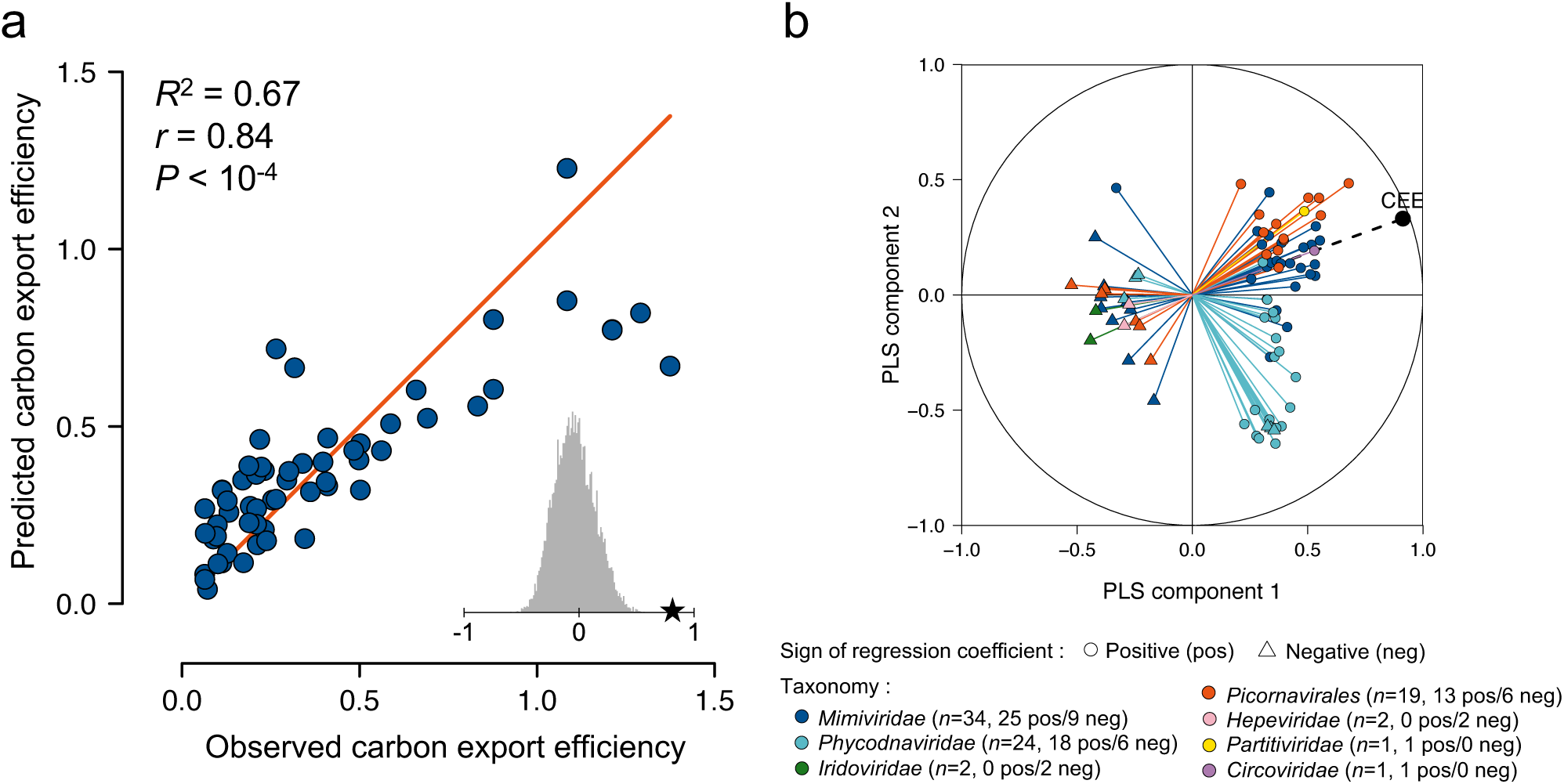
Relative abundance of eukaryotic plankton viruses associated with carbon export efficiency in the global ocean. **a** Bivariate plot between predicted and observed values in a leave-one-out cross-validation test for carbon export efficiency. The PLS regression model was constructed using occurrence profiles of 1,523 marker-gene sequences (1,309 PolBs, 180 RdRPs and 34 Reps) derived from environmental samples. *r*, Pearson correlation coefficient; *R*^2^, the coefficient of determination between measured response values and predicted response values. *R*^2^, which was calculated as 1 – SSE/SST (sum of squares due to error and total) measures how successful the fit is in explaining the variance of the response values. The significance of the association was assessed using a permutation test (*n* = 10,000) (grey histogram in **a**). The red diagonal line shows the theoretical curve for perfect prediction. **b** Pearson correlation coefficients between CEE and occurrence profiles of 83 viruses that have VIP scores > 2 (VIPs) with the first two components in the PLS regression model using all samples. PLS components 1 and 2 explained 83% and 11% of the variance of CEE, respectively. Fifty-eight VIPs had positive regression coefficients in the model (shown with circles), and 25 had negative regression coefficients (shown with triangles).

We also tested for their association with estimates of carbon export flux at 150 meters (CE_150_) and NPP. PLS regressions explained 54% and 64% of the variation in CE_150_ and NPP with Pearson correlation coefficients between observed and predicted values of 0.74 (permutation test, *P* < 1 × 10^−4^) and 0.80 (permutation test, *P* < 1 × 10^−4^), respectively (Figure S6). In these three PLS regression models, 83, 86, and 97 viruses were considered to be key predictors (*i.e.*, Variable Importance in the Projection [VIP] score > 2) of CEE, CE_150_, and NPP, respectively. PLS models for NPP and CE_150_ shared a larger number of predictors (52 viruses) compared to the PLS models for NPP and CEE (seven viruses) (two proportion Z-test, *P* = 4.14 × 10^−12^). Consistent with this observation, CE_150_ was correlated with NPP (Pearson’s *r* = 0.77; parametric test, *P* < 1 × 10^−12^). This result implies that the magnitude of export in the analyzed samples was partly constrained by primary productivity. However, CEE was not correlated with NPP (*r* = 0.16; parametric test, *P* = 0.2) or CE_150_ (*r* = 0.002; parametric test, *P* = 0.99). Thus, as expected, primary productivity was not a major driver for the efficiency of carbon export.

The 83 viruses (5% of the viruses included in our analysis) that were associated with CEE with a VIP score > 2 are considered to be important predictors of CEE in the PLS regression (Figure 3b, Supplemental Data 1), and these viruses are hereafter referred to as VIPs (Viruses Important in the Prediction). Fifty-eight VIPs had positive regression coefficient, and 25 had negative regression coefficient in the prediction (Figure 3b). Most of the positively associated VIPs showed high relative abundance in the Mediterranean Sea and in the Indian Ocean where CEE tends to be high compared with other oceanic regions (Figure 4). Among them, 15 (red labels in Figure 4) also had high relative abundance in samples from other oceanic regions, showing that these viruses are associated with CEE at a global scale. In contrast, negatively associated VIPs tend to have higher relative abundance in the Atlantic Ocean and the Southern Pacific Ocean where CEE is comparatively lower. In the following sections, we investigate potential hosts of the VIPs in order to interpret the statistical association between viral community composition and CEE in the light of previous observations in the literature.

**Figure 4:**
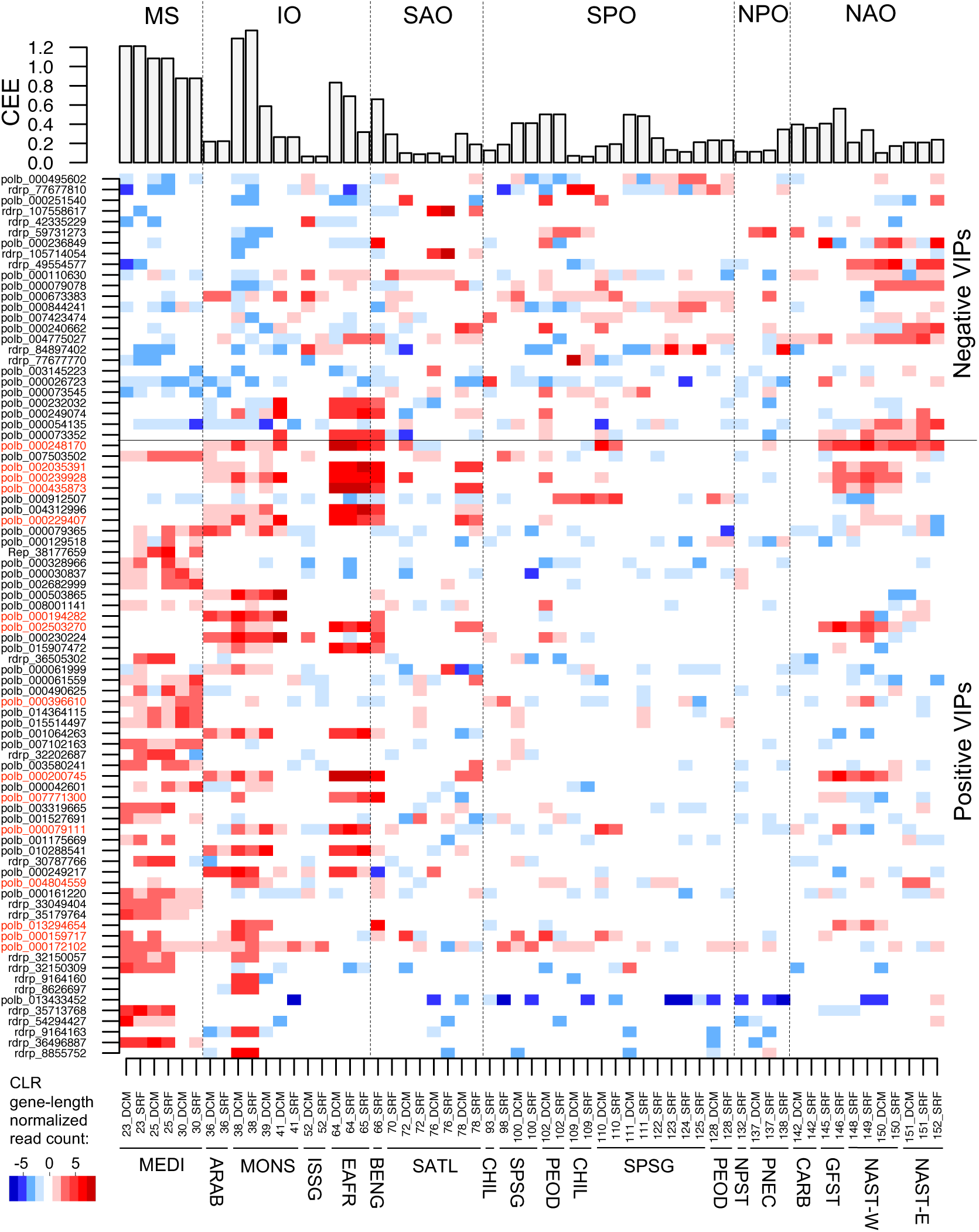
Biogeography of viruses associated with carbon export efficiency. The upper panel shows carbon export efficiency (CEE = CE_deep_/CE_surface_) for 59 sampling sites. The bottom panel is a map reflecting relative abundances, expressed as centered log-ratio transformed, gene-length normalized read counts of viruses positively and negatively associated with CEE that have VIP scores > 2 (VIPs). MS, Mediterranean Sea; IO, Indian Ocean; SAO, South Atlantic Ocean; SPO, South Pacific Ocean; NPO, North Pacific Ocean; NAO, North Atlantic Ocean. The bottom horizontal axis is labeled with *Tara* Oceans station numbers, sampling depth (SRF, surface; DCM, deep chlorophyll maximum), and abbreviations of biogeographic provinces. Viruses labeled in red correspond to positive VIPs that tend more represented in more than one biogeographic province.

### Viruses correlated with CEE infect ecologically important hosts

Most of the VIPs (77 of 83) belong to *Mimiviridae* (*n* = 34 with 25 positive VIPs and nine negative VIPs), *Phycodnaviridae* (*n* = 24 with 18 positive VIPs and six negative VIPs), and ssRNA viruses of the order *Picornavirales* (*n* = 19 with 13 positive VIPs and six negative VIPs) (Figure 3b, Table S1). All the phycodnavirus VIPs were most closely related to prasinoviruses infecting Mamiellales, with amino acid sequence percent identities to reference sequences ranging between 35% and 95%. The six remaining VIPs were two NCLDVs of the family *Iridoviridae* negatively associated with CEE, three RNA viruses (two ssRNA viruses of the family *Hepeviridae* negatively associated with CEE and one dsRNA virus of the family *Partitiviridae* positively associated with CEE), and one ssDNA virus of the family *Circoviridae* positively associated with CEE.

Host information may help understand the relationship between these VIPs and CEE. We performed genomic context analysis for PolB VIPs and phylogeny-guided network-based host prediction for PolB and RdRP to infer putative virus–host relationships (see Transparent Methods).

Taxonomic analysis of genes predicted in 10 metagenome-assembled genomes (MAGs) from the eukaryotic size fractions and 65 genome fragments (contigs) assembled from the prokaryotic size fraction encoding VIP PolBs further confirmed their identity as *Mimiviridae* or *Phycodnaviridae* (Figure S7). The size of MAGs ranged between 30 kbp and 440 kbp with an average of 210 kbp (Table S2). The presence of genes with high sequence similarities to cellular genes in a viral genome is suggestive of a virus–host relationship (Monier et al., 2009; Yoshikawa et al., 2019). Two closely related *Mimiviridae* VIPs, PolB 000079111 (positively associated with CEE) and PolB 000079078 (negatively associated with CEE), were phylogenetically close to the pelagophyte virus *Aureococcus anophagefferens virus* (AaV). One MAG (268 kbp in size) corresponding to PolB 000079111 encoded seven genes showing high similarities to genes from Pelagophyceae, and another MAG (382 kbp in size), corresponding to PolB 000079078, encoded five genes similar to genes from Pelagophyceae. All but one of these 12 genes was encoded on a genome fragment containing genes annotated as viral, including five NCLDV core genes (Supplemental Data 2), excluding the possibility of contamination in these MAGs. Two closely related *Phycodnaviridae* VIPs, PolB 001064263 and 010288541, were positively associated with CEE. Both of these PolBs correspond to a MAG (134 kbp in size) encoding one gene likely derived from Mamiellales. The genomic fragment harboring this cellular gene was found to encode 10 genes annotated as viral (Supplemental Data 2).

We conducted a phylogeny-guided, network-based host prediction analysis for *Mimiviridae*, *Phycodnaviridae*, and *Picornavirales* (Figures S8 and S9). Only a subset of the VIPs was included in this analysis because we kept the most reliable sequences to obtain a well-resolved tree topology. Within the *Prasinovirus* clade, which contained thirteen VIPs (nine positive and four negative), seven different eukaryotic orders were detected as predicted host groups for 10 nodes in the tree. Mamiellales, the only known host group of prasinoviruses, was detected at eight nodes (five of them had no parent-to-child relationships), whereas the other six eukaryotic orders were found at only one node (or two in the case of Eutreptiales) (Figure S8). The order Mamiellales includes three genera (*Micromonas*, *Ostreococcus*, and *Bathycoccus*), which are bacterial-sized green microalgae common in coastal and oceanic environments and are considered to be influential actors in oceanic systems (Monier et al., 2016). Various prasinoviruses (fourteen with available genome sequences) have been isolated from the three genera.

Within the family *Mimiviridae*, which contains fifteen VIPs (10 positive and five negative), twelve different orders were predicted as putative host groups (Figure S8). Collodaria was detected at 15 nodes (two of them had no parent-to-child relationships), and Prymnesiales at six nodes (three of them had no parent-to-child relationships), whereas all other orders were present at a maximum of one node each with no parent-to-child relationships. The nodes enriched for Prymnesiales and Collodaria fell within a monophyletic clade (marked by a red arrow in Figure S8) containing four reference haptophyte viruses infecting Prymnesiales and two reference haptophyte viruses infecting Phaeocystales. Therefore, the environmental PolB sequences in this *Mimiviridae* clade (including five positive VIPs and one negative VIP) are predicted to infect Prymnesiales or related haptophytes. The detection of Collodaria may be the result of indirect associations that reflect a symbiotic relationship with Prymnesiales, as some acantharians, evolutionarily related to the Collodaria, are known to host Prymnesiales species (Mars Brisbin et al., 2018). Known species of Prymnesiales and Phaeocystales have organic scales, except one Prymnesiales species, *Prymnesium neolepis*, which bears siliceous scales (Yoshida et al., 2006). Some species can form blooms and colonies. Previous studies revealed the existence of diverse and abundant noncalcifying picohaptophytes in open oceans (Endo et al., 2018; Liu et al., 2009). Haptophytes as a whole have been estimated to contribute from 30% to 50% of the total photosynthetic standing stock across the world ocean (Hirata et al., 2011; Liu et al., 2009). They constitute an important mixotrophic group in oligotrophic waters (Endo et al., 2018), and mixotrophy is proposed to increase vertical carbon flux by enabling the uptake of organic forms of nutrients (Ward and Follows, 2016). Clear host prediction was not made for the other nine *Mimiviridae* VIPs shown in the phylogenetic tree. Three VIPs (two positive and one negative) in the tree were relatives of AaV. One negatively associated VIP was a relative of *Cafeteria roenbergensis virus* infecting a heterotrophic protist. The five remaining *Mimiviridae* VIPs are very distant from any known *Mimiviridae*.

Sixteen *Picornavirales* VIPs (eleven positive and five negative) were included in the phylogeny-guided, network-based host prediction analysis (Figure 9). Nine (seven positive and two negative) were grouped within *Dicistroviridae* (known to infect insects) and may therefore infect marine arthropods such as copepods, the most ubiquitous and abundant mesozooplankton groups involved in carbon export (Turner, 2015). Three other *Picornavirales* VIPs were placed within a clade containing known bacillarnaviruses. Two of them (35179764 and 33049404) were positively associated with CEE and had diatoms of the order Chaetocerotales as a predicted host group. The third one (107558617) was negatively associated with CEE and distant from other bacillarnaviruses, and had no host prediction. Diatoms have been globally observed in the deep sea (Agusti et al., 2015; Leblanc et al., 2018) and identified as important contributors of the biological carbon pump (Tréguer et al., 2018). One positively associated VIP (32150309) was in a clade containing *Aurantiochytrium single-stranded RNA virus* (AsRNAV), infecting a marine fungoid protist thought to be an important decomposer (Takao et al., 2005). The last three *Picornavirales* VIPs (59731273, 49554577, and 36496887) had no predicted host and were too distant from known *Picornavirales* to speculate about their putative host group.

Outside *Picornavirales*, three RNA virus VIPs (two *Hepeviridae*, negatively associated, and one *Partitiviridae*, positively associated) were identified, for which no reliable host inferences were made by sequence similarity. Known *Hepeviridae* infect metazoans, and known *Partitiviridae* infect fungi and plants. The two *Hepeviridae*-like viruses were most closely related to viruses identified in the transcriptomes of mollusks (amino acid identities of 48% for 42335229 and 43% for 77677770) (Shi et al., 2016). The *Partitiviridae*-like VIP (35713768) was most closely related to a fungal virus, *Penicillium stoloniferum virus S* (49% amino acid identity).

One ssDNA virus VIP (38177659) was positively associated with CEE. It was annotated as a *Circoviridae*, although it groups with other environmental sequences as an outgroup of known *Circoviridae*. This VIP was connected with copepod, mollusk, and Collodaria OTUs in the co-occurrence network but no enrichment of predicted host groups was detected for its clade. *Circoviridae*-like viruses are known to infect copepods (Dunlap et al., 2013) and have been reported to associate with mollusks (Dayaram et al., 2015), but none have been reported for Collodaria.

Overall, we could infer hosts for 36 VIPs (Tables S3 and S4). Most of the predicted hosts are known to be ecologically important as primary producers (Mamiellales, Prymnesiales, Pelagophyceae, and diatoms) or grazers (copepods). Of these, diatoms and copepods are well known as important contributors to the BCP but others (*i.e.*, Mamiellales, Prymnesiales, Pelagophyceae) have not been recognized as major contributors to the BCP. Our analysis also revealed that positive and negative VIPs are not separated in either the viral or host phylogenies.

### Viruses positively correlated with CEE tend to interact with silicified organisms

The phylogeny-guided, network-based host prediction analysis correctly predicted known virus–host relationships (for viruses infecting Mamiellales, Prymnesiales, and Chaetocerotales) using our large dataset, despite the reported limitations of these co-occurrence network-based approaches (Coenen and Weitz, 2018). This result prompted us to further exploit the species co-occurrence networks (Table S5) to investigate functional differences between the eukaryotic organisms predicted to interact with positive VIPs, negative VIPs, and viruses less important for prediction of CEE (VIP score < 2) (non-VIPs). Positive VIPs had a greater proportion of connections with silicified eukaryotes (*Q* = 0.001), but not with chloroplast-bearing eukaryotes (*Q* = 0.16) nor calcifying eukaryotes (*Q* = 1), compared to non-VIPs (Table S6). No functional differences were observed between negative VIPs and non-VIPs viruses (Table S6) or positive VIPs (Table S7).

### Multifarious ways viruses affect the fate of carbon

Our analysis revealed that eukaryotic virus composition was able to predict CEE in the global sunlit ocean and 83 out of the 1,523 viruses had a high importance in the predictive model. This association is not a proof that the viruses are the cause of the variation of CEE. For example, a virus may be found to be associated with CEE if its host affects CEE regardless of viral infection. This would be the case especially if latent/persistent viruses are widespread and abundant in phytoplankton (Goic and Saleh, 2012). Organisms that preferentially grow in marine snow (Bochdansky et al., 2017) may bring associations between viruses infecting those organisms and CEE; this could be the case for the AsRNAV-related VIP that we identified. Alternatively, the observed associations between VIPs and CEE may reflect a more direct causal relationship, which we attempt to explore in light of the large body of literature on the mechanisms by which viruses impact the fate of carbon in the oceans.

Among the 83 VIPs, 58 were positively associated with CEE. Such a positive association is expected from the “viral shuttle” model, which states that viral activity could facilitate carbon export to the deep ocean (Fuhrman, 1999; Sullivan et al., 2017; Weinbauer, 2004) because viral infection can facilitate cell sinking (Lawrence and Suttle, 2004) and increase the sizes of particles (Peduzzi and Weinbauer, 1993; Yamada et al., 2018); for instance, a virus may induce secretion of sticky material that contributes to cell/particle aggregation, such as transparent exopolymeric particles (TEP) (Nissimov et al., 2018). The data we used to estimate carbon export are based on the particle size distribution and concentration, and do not convey information regarding the aggregation status of particles. Therefore, we cannot directly test for a relation between viruses and aggregation at the sampling sites. Nonetheless, we found that CEE (*i.e.*, CE_deep_/CE_surface_) increased with the change of particles size from surface to deep (*ρ* = 0.42, *P* = 8 × 10^−9^) (Figure S10). This positive correlation may reflect an elevated level of aggregation (either enhanced by viral activity or not) in places where CEE is high, although it could be also due to the presence of large organisms at depth.

Greater aggregate sinking along with higher particulate carbon fluxes was observed in North Atlantic blooms of *Emiliania huxleyi* that were infected early by the virus EhV, compared with late-infected blooms (Laber et al., 2018). In the same bloom, viral infection stage was found to proceed with water column depth (Sheyn et al., 2018). Enhanced TEP production for these same early infected calcifying populations was observed over a three-day period in deck-board bottle incubations (Laber et al., 2018). Laboratory observations also exist for enhanced TEP production and aggregate formation during the early phase of EhV infection of a calcifying *E. huxleyi* strain (Laber et al., 2018; Nissimov et al., 2018). These observations strongly suggest that infection-induced TEP production in organisms containing dense material (*e.g.*, calcite scales for *E. huxleyi*) can facilitate carbon export. No EhV-like PolB sequences were detected in our dataset, which was probably due to sampled areas and seasons.

Laboratory experiments suggest that viruses closely related to positive VIPs, such as prasinoviruses, have infectious properties that may drive carbon export. Cultures of *Micromonas pusilla* infected with prasinoviruses showed increased TEP production compared with non-infected cultures (Lønborg et al., 2013), although it is not known if this increase leads to aggregation. The hosts of prasinoviruses have been proposed to contribute to carbon export because they were observed in abyssopelagic zone at sampling sites dominated by Mamiellales in their surface waters in the western subtropical North Pacific (Shiozaki et al., 2019). Some prasinoviruses encode glycosyltransferases (GTs) of the GT2 family. Similar to the a098r gene (GT2) in *Paramecium bursaria Chlorella virus 1*, the expression of GT2 family members during infection possibly leads to the production of a dense fibrous hyaluronan network at the surface of infected cells. Such a network may trigger the aggregation of host cells, facilitate viral propagation (Van Etten et al., 2017), and increase the cell wall C:N ratio. We detected one GT2 in a MAG of two *Phycodnaviridae*-like positive VIPs (000200745 and 002503270) predicted to infect Mamiellales, one in a MAG corresponding to the putative pelagophyte positive VIP 000079111 related to AaV and six in two MAGs (three each) corresponding to two *Mimiviridae*-like positive VIPs (000328966 and 001175669). *Phaeocystis globosa virus* (PgV), closely related to the positive VIP PolB 000912507 (Figure S8), has been linked with increased TEP production and aggregate formation during the termination of a *Phaeocystis* bloom (Brussaard et al., 2007). Two closely related bacillarnavirus VIPs were positively associated with CEE and predicted to infect Chaetocerales. A previous study revealed an increase in abundance of viruses infecting diatoms of *Chaetoceros* in both the water columns and the sediments during the bloom of their hosts in a coastal area (Tomaru et al., 2011), suggesting sinking of cells caused by viruses. Furthermore, the diatom *Chaetoceros tenuissimus* infected with a DNA virus (CtenDNAV type II) has been shown to produce higher levels of large-sized particles (50 to 400 μm) compared with non-infected cultures (Yamada et al., 2018).

The other 25 VIPs were negatively associated with CEE. This association is compatible with the “viral shunt,” which increases the amount of DOC (Wilhelm and Suttle, 1999) and reduces the transfer of carbon to higher trophic levels and to the deep ocean (Fuhrman, 1999; Weitz et al., 2015). Increased DOC has been observed in culture of Mamiellales lysed by prasinoviruses (Lønborg et al., 2013). Although this culture-based observation may be difficult to extrapolate to natural conditions, where the cell concentration and thus the contact rate with viruses are probably lower, Mamiellales species are known to form blooms during which cell densities may be comparable with cultures (Zhu et al., 2005). A field study reported that PgV, to which the negative VIP PolB 000054135 is closely related (Figure S8), can be responsible for up to 35% of cell lysis per day during bloom of its host (Baudoux et al., 2006), which is likely accompanied by consequent DOC release. Similarly, the decline of a bloom of the pelagophyte *Aureococcus anophagefferens* has been associated with active infection by AaV (to which one negative VIP is closely related) (Moniruzzaman et al., 2017). Among RNA viruses, eight were negative VIPs (six *Picornavirales* and two *Hepeviridae*). The higher representation of *Picornavirales* in the virioplankton (Culley, 2018) than within cells (Urayama et al., 2018) suggests that they are predominantly lytic, although no information exists regarding the effect of *Picornavirales* on DOC release.

It is likely that the “viral shunt” and “viral shuttle” simultaneously affect and modulate CEE in the global ocean (Zimmerman et al., 2019a). The relative importance of these two phenomena must fluctuate considerably depending on the host traits, viral effects on metabolism, and environmental conditions. Reflecting this complexity, viruses of a same host group could be found to be either positively or negatively associated with CEE. For example, among prasinoviruses most likely infecting Mamiellales, 18 were positive VIPs and six were negative VIPs. Two closely related prasinoviruses (sharing 97.5% genome-wide identity) are known to exhibit different ecological strategies with notably distinct molecular signatures on the organic matter released upon infection of the same host (Zimmerman et al., 2019b). We found that even two very closely related *Mimiviridae* viruses (PolBs 000079111 and 000079078 sharing 94% nucleotide identity over their full gene lengths) most likely infecting pelagophyte algae were positively and negatively associated with CEE. Furthermore, it is known that an early-infected *E. huxleyi* system was linked with both higher aggregation “at surface” and higher remineralization “at deep” compared to late-infected blooms (Laber et al., 2018). Therefore, the viral effect on carbon cycle may vary also with depth.

Five percent of the tested viruses were associated with CEE in our study. Similarly, four percent of bacterial virus populations were found to be associated with the magnitude of carbon export at 150 meters (Guidi et al., 2016). These results suggest that viruses affecting carbon export are rather uncommon. It is plausible that such viruses affect CEE by infecting organisms that are functionally important (abundant or keystone species), as we observed in host prediction. The vast majority (95%) of non-VIPs may not have a significant impact on CEE, because they do not strongly impact the host population, for instance, by stably coexisting with their hosts. It is worth noting that experimental studies have reported cultures of algae with viruses that reach a stable co-existence state after a few generations (Yau et al., 2020). It is also possible that some of these non-VIPs can impact carbon export but were not captured in the infection stage affecting the export process. Viruses captured in our samples can represent active viruses in different infectious stages (early, mid or late) for metagenomes and metatranscriptomes or at the post-lysis stage for metagenomes.

### Potential effects of global climate changes on viral shunt/shuttle and CEE

Increasing evidence suggests that the biological carbon pump is highly dependent on the planktonic community composition, and as discussed above, viruses represent a possible key parameter that determines the efficiency of carbon export. In the photic layer of the oceans, the composition of planktonic communities is strongly affected by sea surface temperature (Salazar et al., 2019; Sunagawa et al., 2015), and CEE may therefore be affected by ocean warming. Our result indicated that viruses infecting small phytoplankton such as Mamiellales and haptophytes are likely associated with CEE. Interestingly, many studies showed that high temperature and/or CO_2_ levels are associated with an increased contribution of small sized phytoplankton to the total biomass (Hare et al., 2007; Mousing et al., 2014; Sugie et al., 2020).

An increase in CO_2_ level in the surface seawater also causes a decrease in pH (i.e., ocean acidification). Previous studies demonstrated that the decrease in seawater pH negatively affect the growth of calcified and silicified phytoplankton cells (Doney et al., 2009; Endo et al., 2016; Petrou et al., 2019). The biogenic minerals such as calcium carbonate and silica act as ballasts in sinking particles (Iversen and Ploug, 2010). Given the statistical association that we detected between the viruses positively correlated with CEE and the silicified predicted host planktons, the ocean acidification may decrease the viral shuttle and thus CEE globally in the future.

The increased sea surface temperature will decrease the nutrient supply at the surface of the oligotrophic ocean by preventing the vertical mixing. The decrease in nutrient availability of surface seawaters possibly diminishes the net primary production (NPP) and the magnitude of carbon export (corresponded to CE_150_ in our study) (Riebesell et al., 2009). Consistently, a downward trend of global phytoplankton abundance has been observed by satellite (Boyce et al., 2010). In such a scenario of the global decrease of NPP, the efficiency of export would be an important factor for a precise estimation of carbon export in the future ocean. In this regard, the role of marine viruses in the carbon cycle and export should be further investigated and eventually be integrated into prospective models for the climate change.

## Conclusions

Eukaryotic virus community composition was able to predict CEE at 59 sampling sites in the photic zone of the world ocean. This statistical association was detected based on a large omics dataset collected throughout the oceans and processed with standardized protocols. The predictability of CEE by viral composition is consistent with the hypothesis that “viral shuttle” and “shunt” are functioning at a global scale. Among 83 viruses with a high importance in the prediction of CEE, 58 viruses were positively and 25 negatively correlated with carbon export efficiency. Most of these viruses belong to *Prasinovirus*, *Mimiviridae*, and *Picornavirales* and are either new to science or with no known roles in carbon export efficiency. Thirty-six of these “select” viruses were predicted to infect ecologically important hosts such as green algae of the order Mamiellales, haptophytes, diatoms, and copepods. Positively associated viruses had more predicted interactions with silicified eukaryotes than non-associated viruses did. Overall, these results imply that the effect of viruses on the “shuttle” and “shunt” processes could be dependent on viral hosts and ecosystem dynamics.

## Limitations of the study

The observed statistical associations between viral compositions and examined parameters (*i.e.*, CEE, CE and NPP) do not convey the information about the direction of their potential causality relationships, and they could even result from indirect relationships as discussed above. Certain groups of viruses detected in samples may be over- or under-represented because of the technical limitations in size fractionation, DNA/RNA extraction and sequencing.

## Resource Availability

### Lead Contact

Further information and requests for resources and reagents should be directed to and will be fulfilled by Lead Contact, Hiroyuki Ogata.

### Materials Availability

This study did not generate unique reagent.

### Data and Code Availability

The authors declare that the data supporting the findings of this study are available within the paper and its supplemental files, as well as at the GenomeNet FTP: ftp://ftp.genome.jp/pub/db/community/tara/Cpump/Supplementary_material/.

Our custom R script used to test for association between viruses and environmental variables (CEE, CE_150_ and NPP) is available along with input data at the GenomeNet FTP: ftp://ftp.genome.jp/pub/db/community/tara/Cpump/Supplementary_material/PLSreg/.

The Taxon Interaction Mapper (TIM) tool developed for this study and used for virus host prediction is available at https://github.com/RomainBlancMathieu/TIM.

## Supplemental Files

- Supplemental_Information.pdf: supplemental figures and tables, and transparent methods
- Supplemental_Data_1_2.xlsx

## Acknowledgements

We thank the *Tara* Oceans consortium, the projects Oceanomics and France Genomique (grants ANR-11-BTBR-0008 and ANR-10-INBS-09), and the people and sponsors who supported the *Tara* Oceans Expedition (http://www.embl.de/tara-oceans/) for making the data accessible. This is contribution number XXX of the *Tara* Oceans Expedition 2009–2013. Computational time was provided by the SuperComputer System, Institute for Chemical Research, Kyoto University. We thank Barbara Goodson, Ph.D., and Sara J. Mason, M.Sc., from Edanz Group (https://en-author-services.edanzgroup.com/) for editing a draft of this manuscript. This work was supported by JSPS/KAKENHI (Nos. 26430184, 18H02279, and 19H05667 to H.O. and Nos. 19K15895 and 19H04263 to H.E.), Scientific Research on Innovative Areas from the Ministry of Education, Culture, Science, Sports and Technology (MEXT) of Japan (Nos. 16H06429, 16K21723, and 16H06437 to H.O.), the Collaborative Research Program of the Institute for Chemical Research, Kyoto University (2019-29 to S.C.), the Future Development Funding Program of the Kyoto University Research Coordination Alliance (to R.B.M.), the ICR-KU International Short-term Exchange Program for Young Researchers (to S.C.), and the Research Unit for Development of Global Sustainability (to H.O. and T.O.D.).

## Author contributions

H.O. and R.B.M. conceived the study. R.B.M and H.K. performed most of the analyses. H.E. and L.G. designed carbon export analysis. R.H.V and S.C. performed network analysis. N.H. and C.d.V. analyzed eukaryotic sequences. T.O.D., M.G., P.F. and O.J. analyzed viral MAGs. C.H.N. and H.M. contributed to statistical analysis. M.B.S. and C.A.S. contributed to interpretations. All authors edited and approved the final version of the manuscript.

## Declaration of Interests

The authors declare no competing interests.

## Supplemental Information

### Supplemental Figures

**Figure S1:**
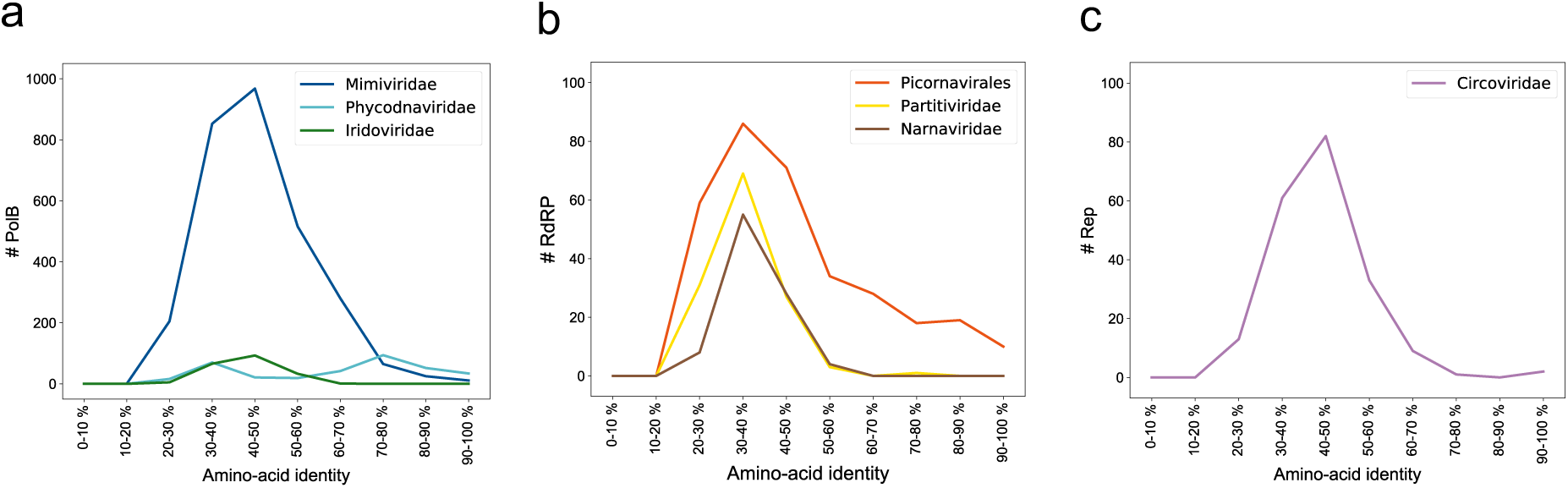
Distribution of the degree (%) of amino acid identity between environmental sequences and their best BLAST hits to reference sequences for nucleocytoplasmic large DNA viruses (NCLDVs) (**a**), RNA viruses (**b**), and ssDNA viruses (**c**).

**Figure S2:**
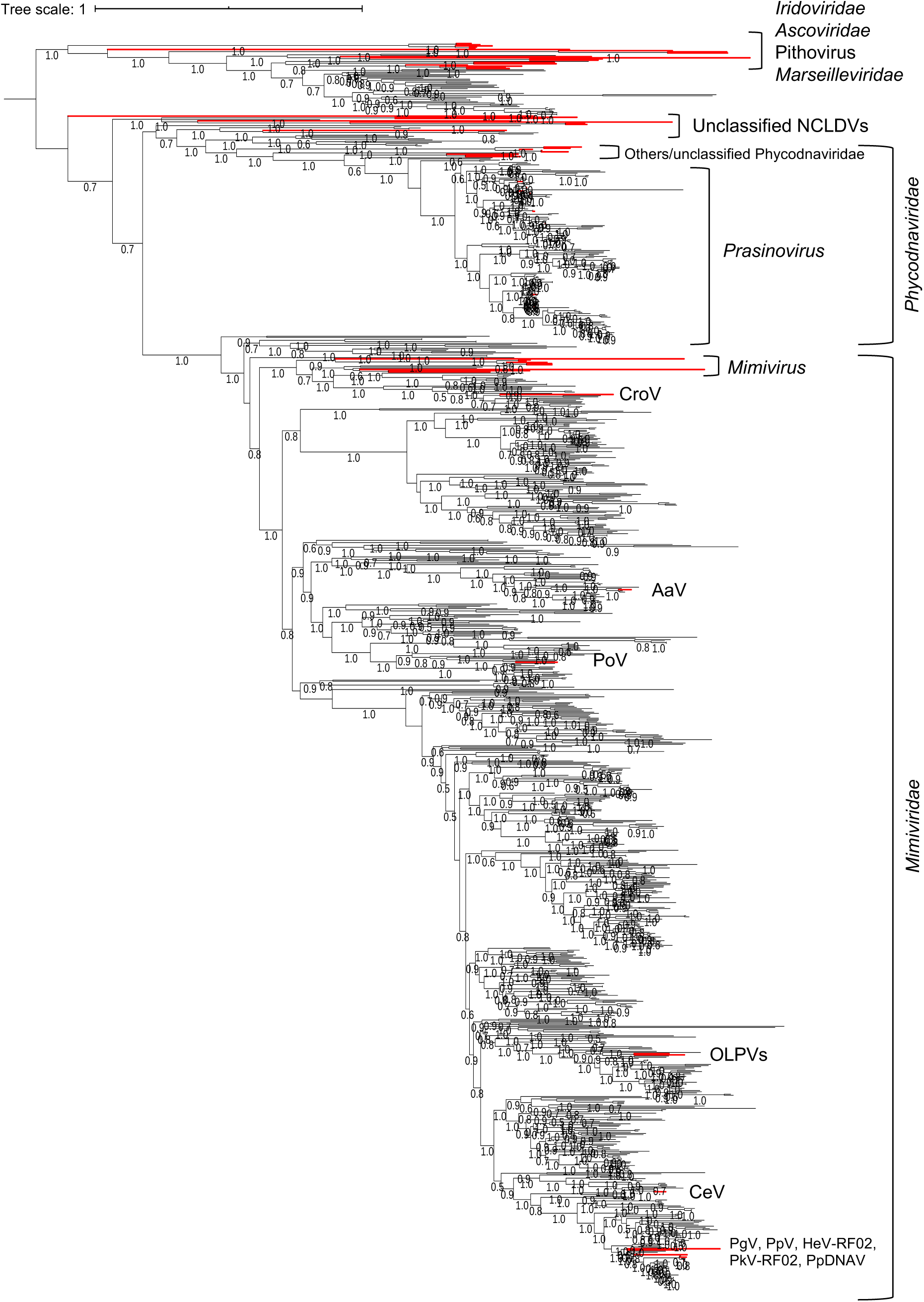
Maximum likelihood phylogenetic trees for NCLDV DNA polymerase family B. Environmental sequences are shown in black and references in red. Approximate Shimodaira– Hasegawa (SH)-like local support values greater than 0.8 are shown. Scale bar indicates one change per site.

**Figure S3:**
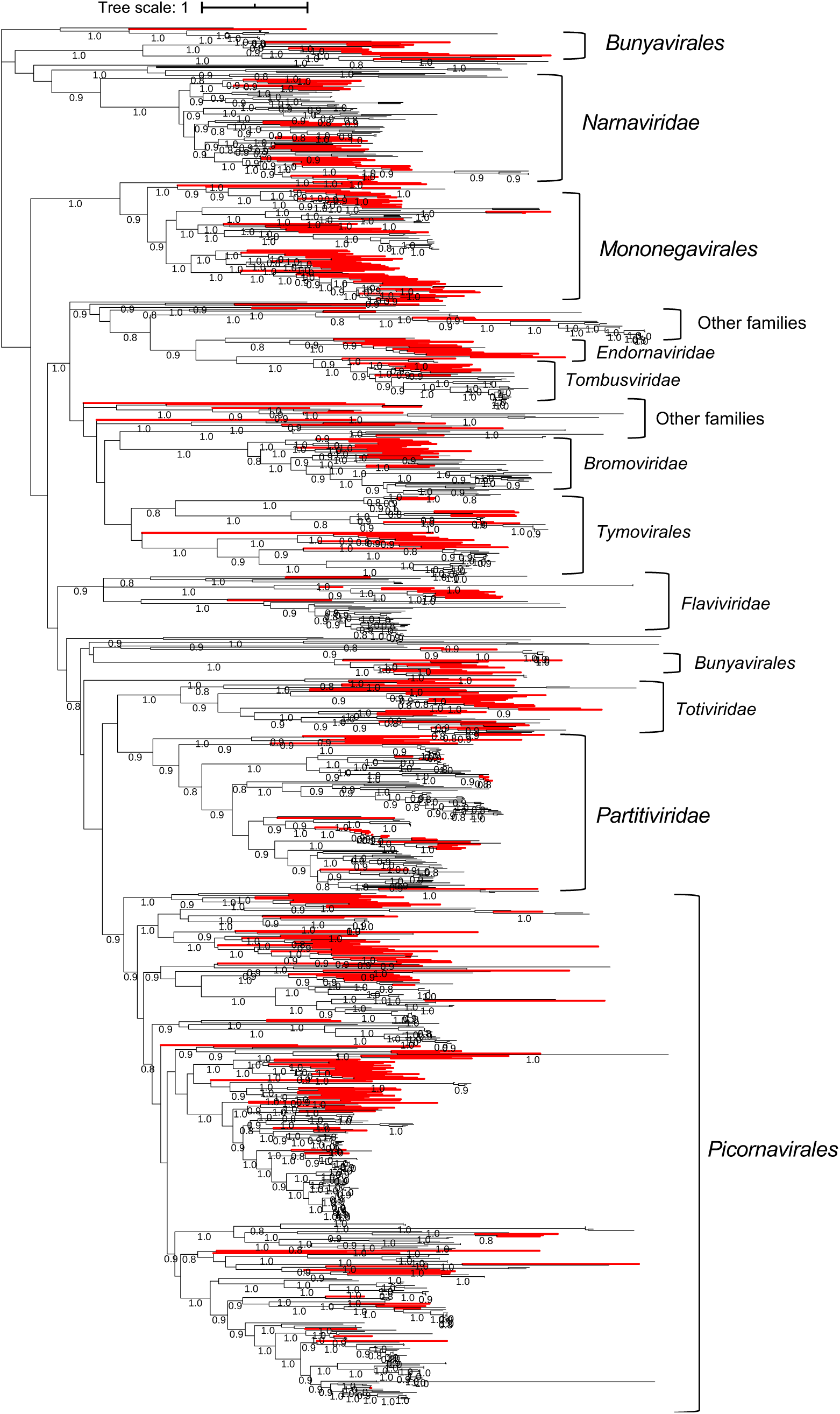
Unrooted maximum likelihood phylogenetic trees for RNA virus RNA-dependent RNA polymerase. Environmental sequences are shown in black and references in red. Approximate SH-like local support values greater than 0.8 are shown. Scale bar indicates one change per site.

**Figure S4:**
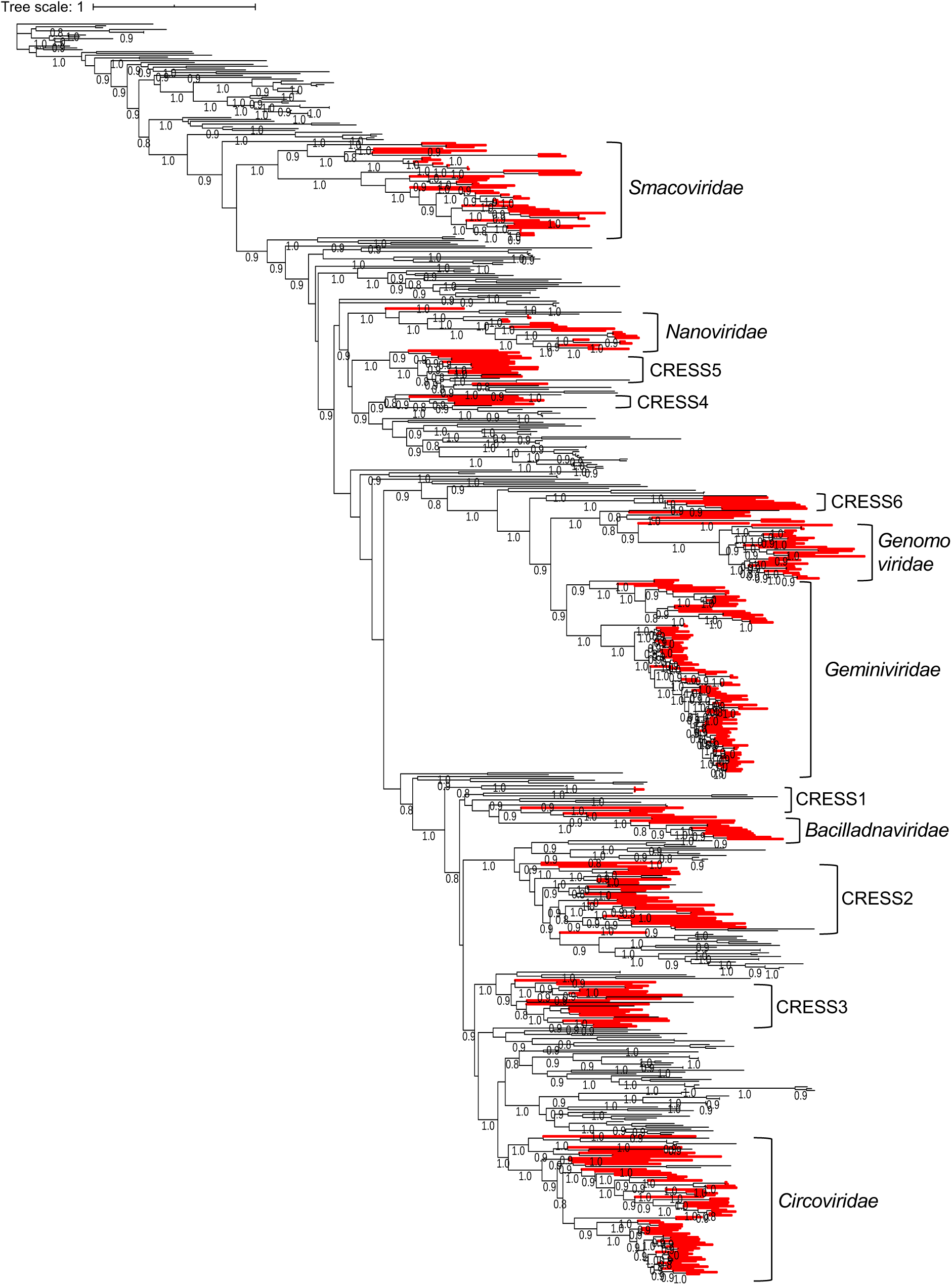
Unrooted maximum likelihood phylogenetic trees for ssDNA virus replication-associated protein. Environmental sequences are shown in black and references in red. Approximate SH-like local support values greater than 0.8 are shown. Scale bar indicates one change per site.

**Figure S5:**
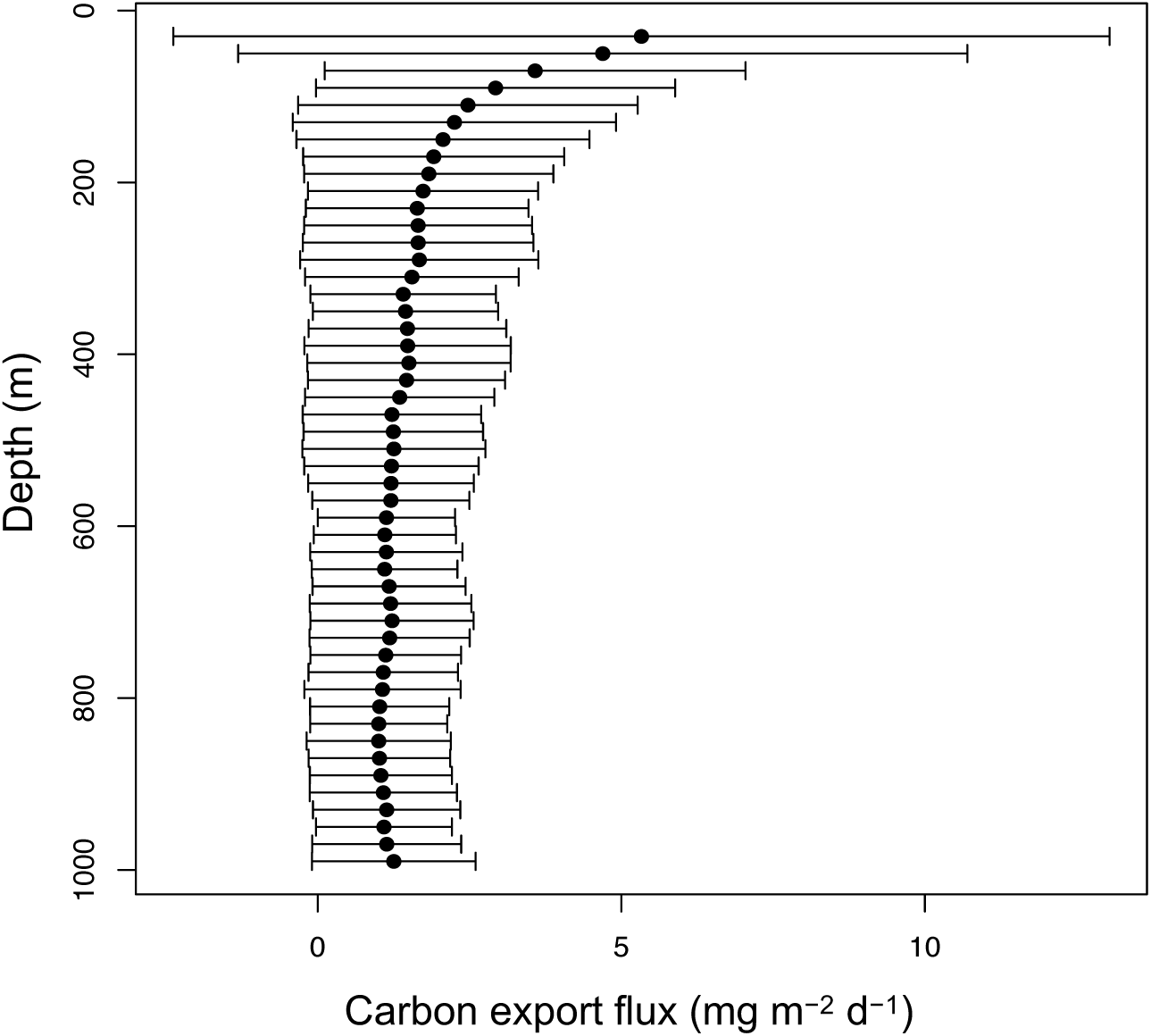
Variation in carbon export flux (mg m^−2^ d^−1^) across sampling depths in the water column. Dots are average values, and horizontal lines represent standard deviations.

**Figure S6:**
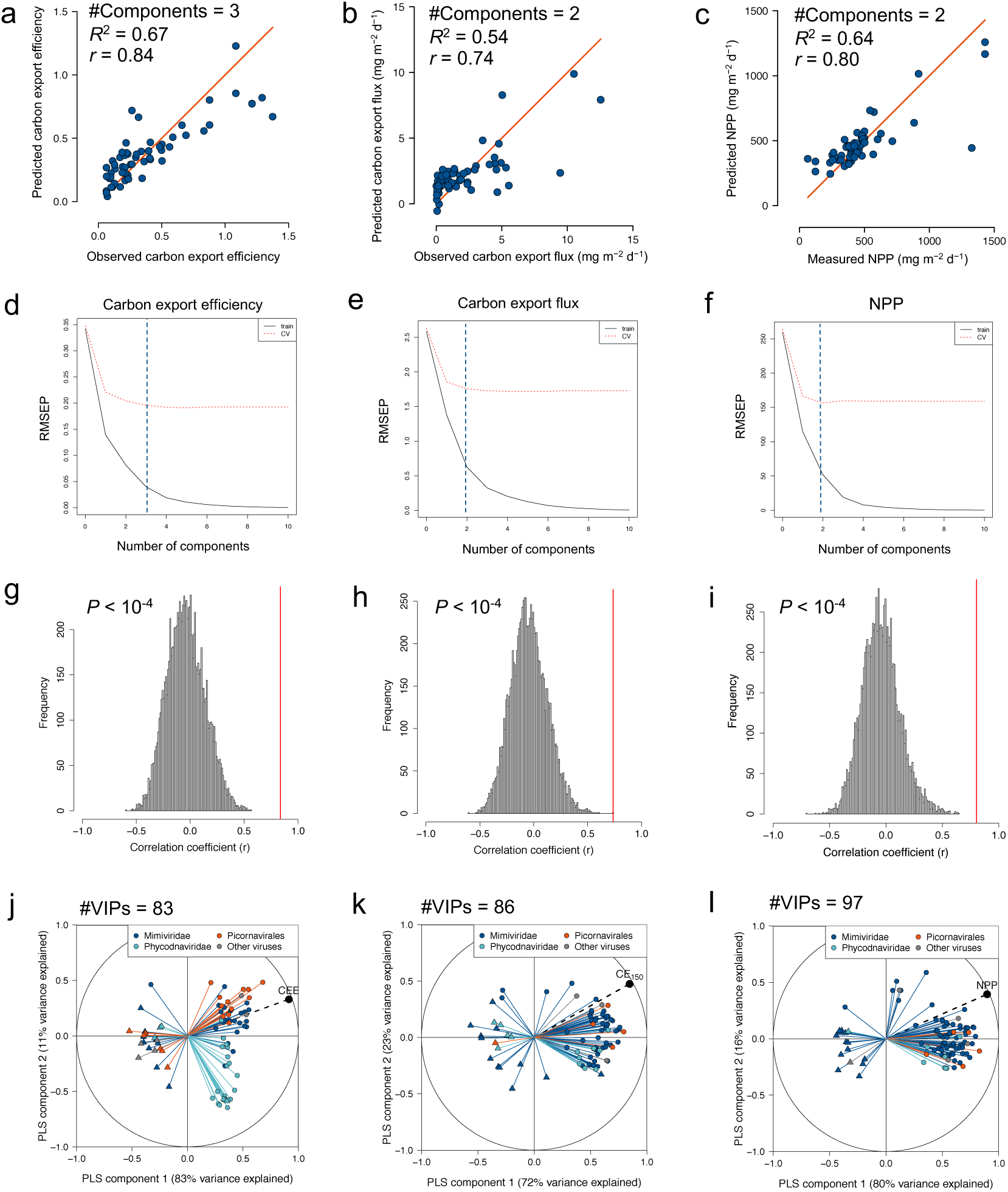
The results of PLS regressions using relative abundance profiles of viral marker-genes to explain the variance of carbon export efficiency (CEE) (**a, d, g, j**), carbon export flux at 150 meters (CE_150_) (**b, e, h, k**), and net primary production (NPP) (**c, f, i, l**). **a–c** Bivariate plots between predicted and observed response values in a leave-one-out cross-validation test. The red diagonal line shows the theoretical curve for perfect prediction. **d–f** Variation in root mean squared error of predictions (RMSEP) for the training set (solid black line) and cross-validation set (red dashed line) across the number of components. Blue dashed line shows the number of components selected for the analysis. **g–i** Results of the permutation tests (*n* = 10,000) supporting the significance of the association between viruses and the response variable. The histograms show the distribution of Pearson correlation coefficients obtained from PLS models reconstructed based on the permutated response variable and red line show the non-permutated response variable. **j–l** Pearson correlation coefficients between the response variable and abundance profiles of viruses with VIP scores > 2 (VIPs) with the first two components in the PLS regression model using all samples. Viruses with positive regression coefficients are shown with circles, and those with negative coefficients are shown with triangles.

**Figure S7:**
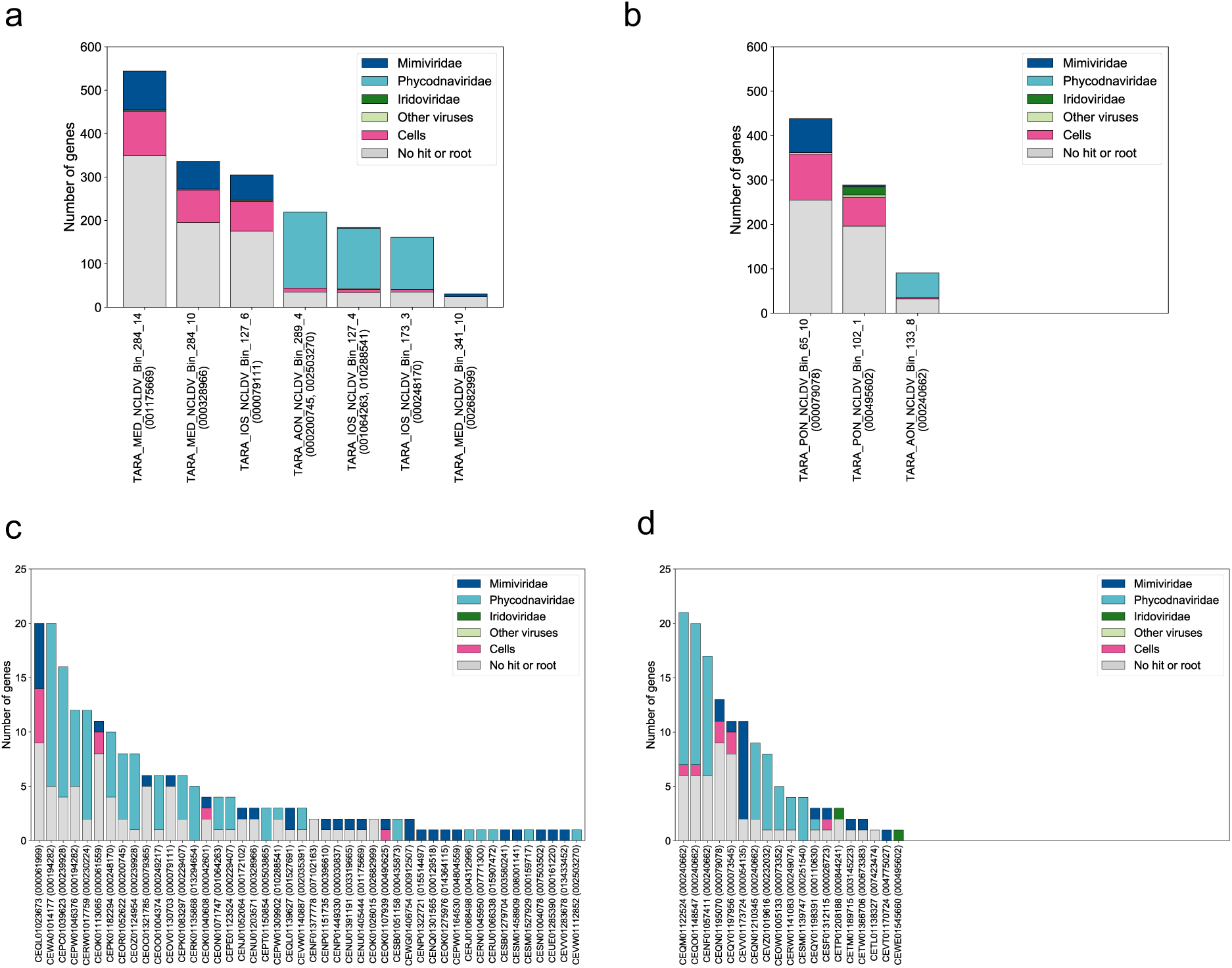
Taxonomic composition of genes predicted in viral genome fragments encoding NCLDV PolBs positively (a and c) or negatively (b and d) associated with CEE (VIP score > 2). **a and b** Metagenome-assembled genomes (MAGs) derived from samples filtered to retain particles of sizes > 0.8 μm. **c and d** Contigs derived from samples filtered to retain particles between 0.2 μm and 3 μm in size. Taxonomic annotations were performed as described in Methods.

**Figure S8:**
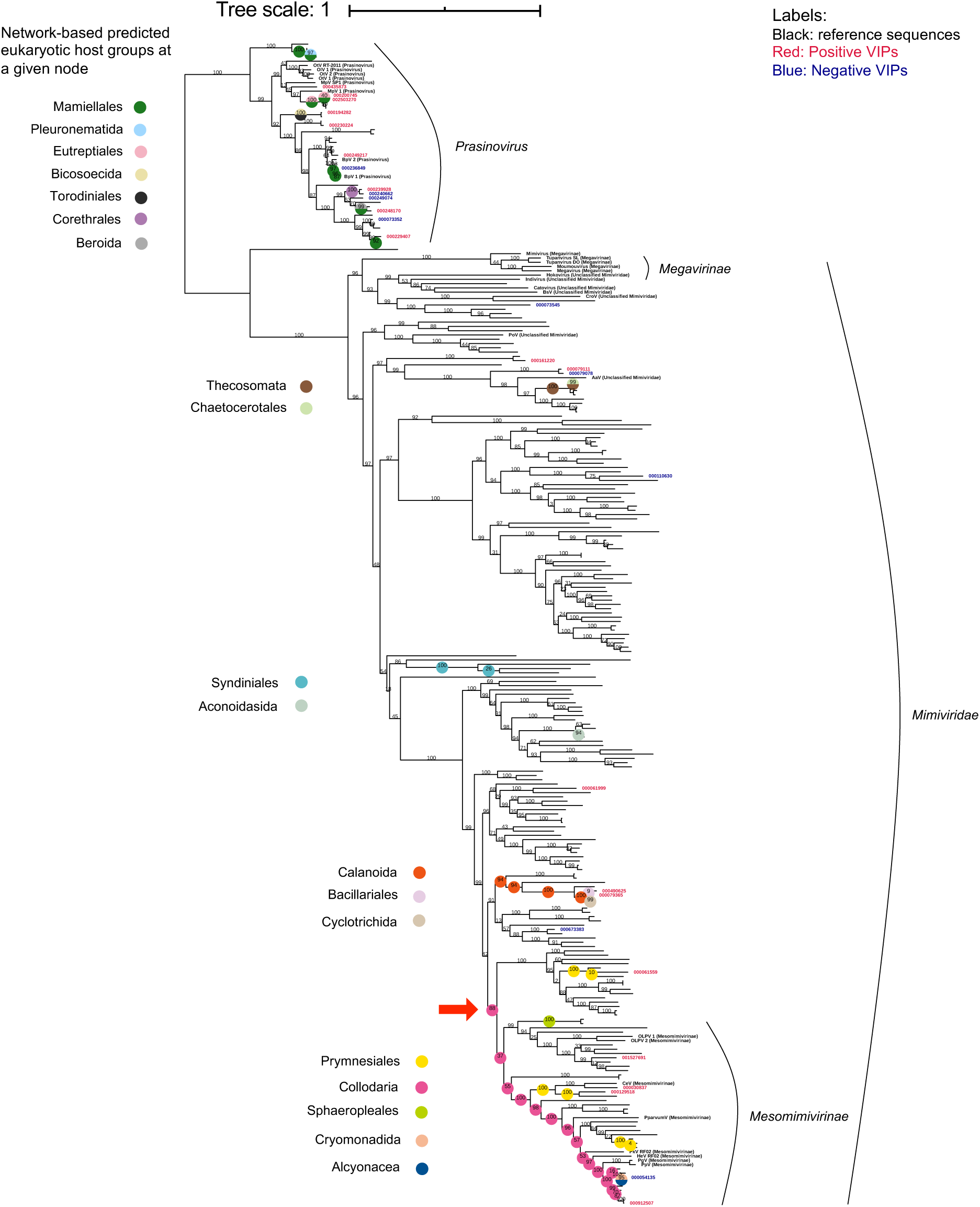
Phylogenetic positions of NCLDV PolBs associated with CEE and network-based predicted eukaryotic host groups. The unrooted maximum likelihood phylogenetic tree contains environmental (labeled in red if VIP score > 2 and the regression coefficient is positive, labeled in blue if negative) and reference (labeled in black) sequences of *Prasinovirus* and *Mimiviridae* PolBs. The approximate SH-like local support values are shown in percentages at nodes, and the scale bar indicates one change per site. Host groups predicted at nodes are shown with colored circles. The red arrow points to a clade of viruses predicted to infect Prymnesiales.

**Figure S9:**
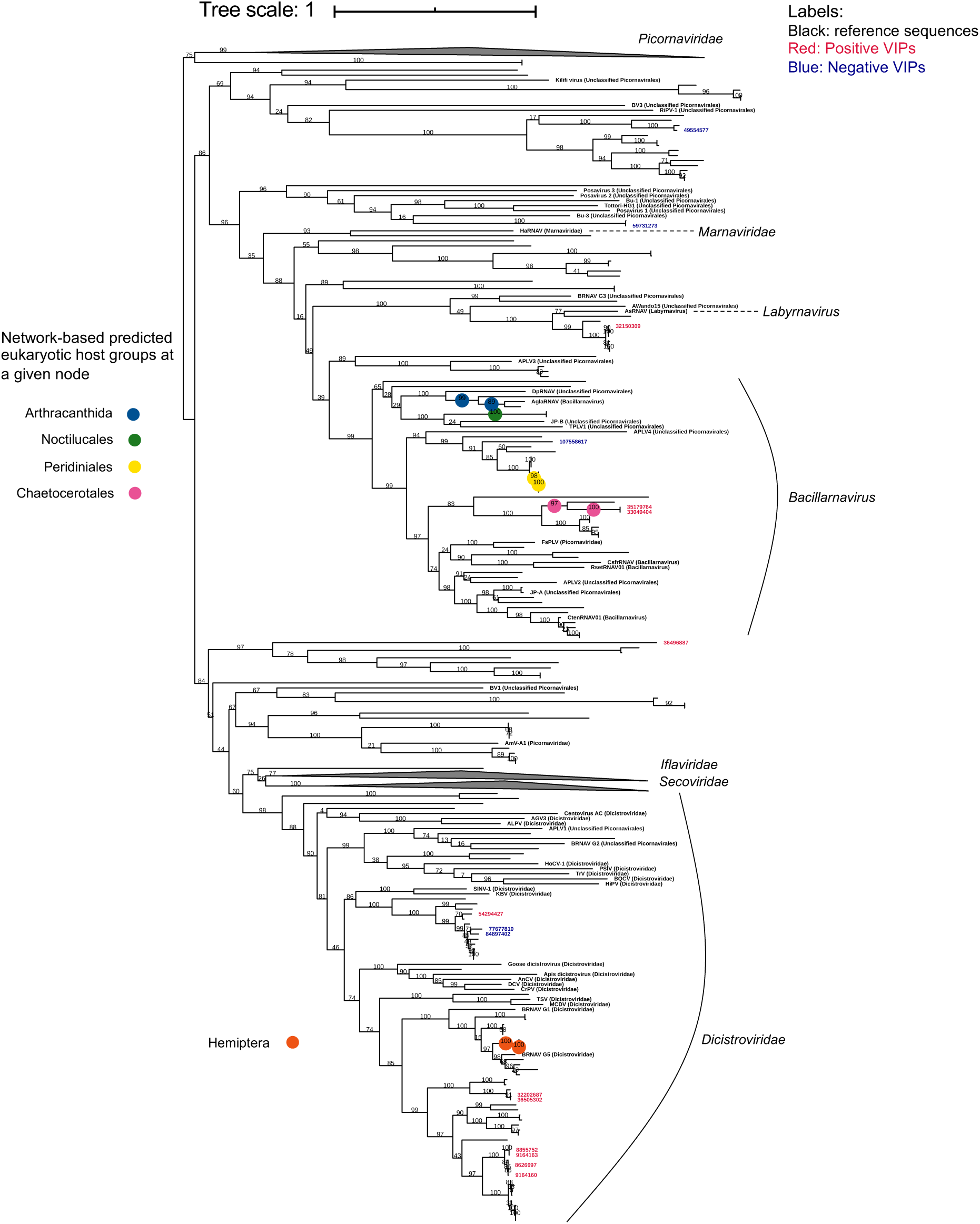
Phylogenetic position of *Piconavirale*s RdRPs associated with CEE and network-based predicted eukaryotic host groups. The unrooted maximum likelihood phylogenetic tree contains environmental (labeled in red if VIP score > 2 and the regression coefficient is positive, labeled in blue if negative) and reference (labeled in black) sequences of *Piconavirale*s RdRPs. The approximate SH-like local support values are shown in percentages at nodes, and the scale bar indicates one change per site. Host groups predicted at nodes are shown with colored circles.

**Figure S10:**
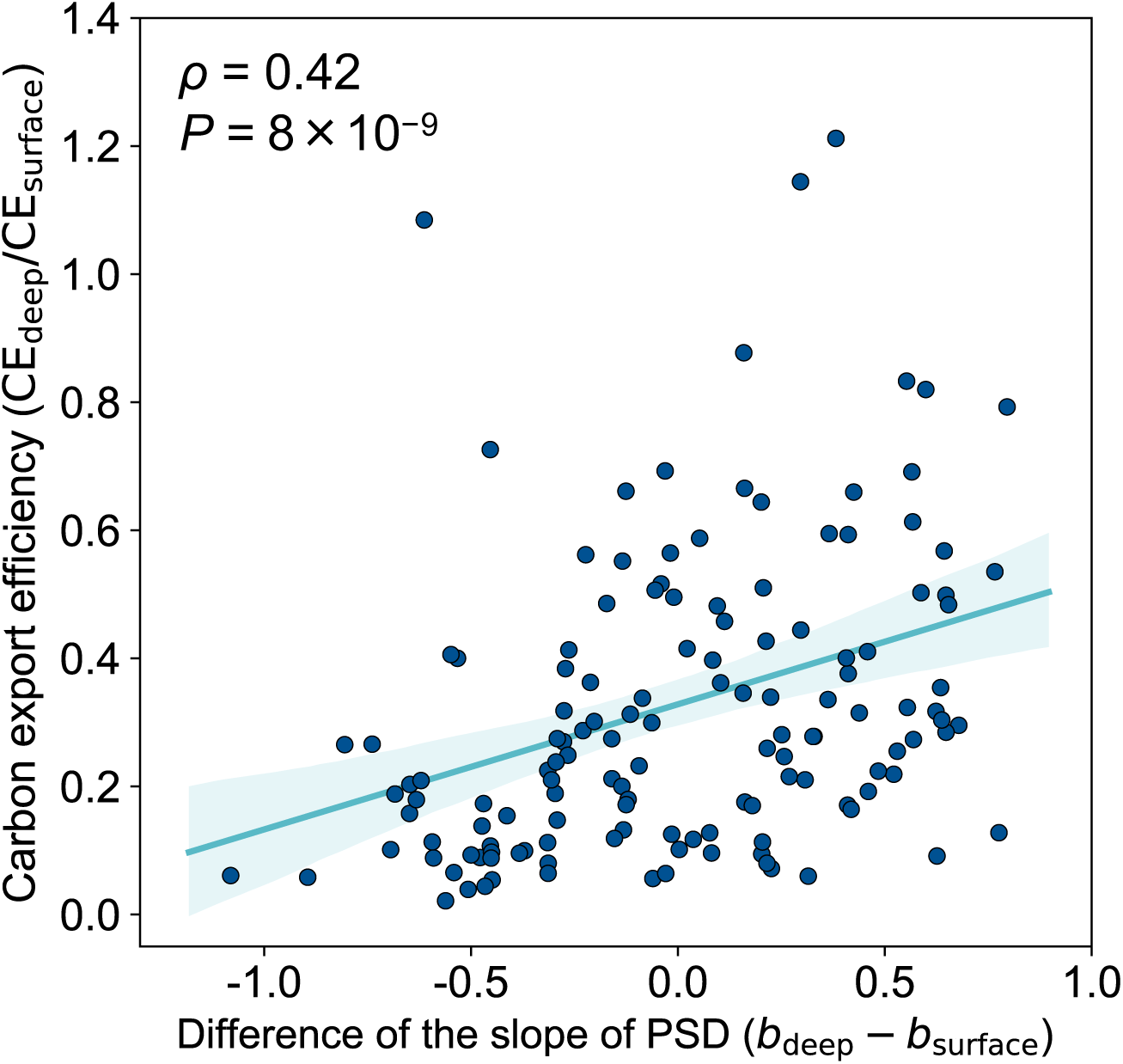
Carbon export efficiency (CEE) is correlated with the change in the slope of particle size distribution (PSD) that occurred from the surface to deep (below the euphotic zone). Observed PSDs were fitted in the form *n* = *ad^b^*, where *n* is the frequency of particles of a given size, *d* is the particle diameter, and *a* and *b* are parameters (as described by(Guidi et al., 2008)). *b*, the PSD slope, is a proxy for particles size. For example *b* = -5 indicates presence of a large proportion of smaller particles, whereas *b* = -3 indicates a preponderance of larger particles. A higher *b* value at deep compared to surface is suggestive of aggregation or presence of larger organisms at deep compare to surface. The blue line shows the regression line between CEE and the PSD slope difference between surface and deep. The shade around the regression line shows the 95% confidence interval.

### Supplemental Tables

**Table S1:** Viral lineages associated with CEE

| Viruses |  | VIPs | Positives<br>VIPs | Negative<br>VIPs |
| --- | --- | --- | --- | --- |
| NCLDV's | Mimiviridae | 34 | 25 | 9 |
|  | Phycodnaviridae | 24 | 18 | 6 |
|  | Iridoviridae | 2 | 0 | 2 |
|  | Other NCLDV's * | 0 | 0 | 0 |
|  | Total | 60 | 43 | 17 |
| RNA viruses | Picornavirales (ssRNA+) | 19 | 13 | 6 |
|  | Partitiviridae (dsRNA) | 1 | 1 | 0 |
|  | Narnaviridae (ssRNA+) | 0 | 0 | 0 |
|  | Other families | 2* | 0 | 2 |
|  | Unclassified | 0 | 0 | 0 |
|  | RNA viruses | 0 | 0 | 0 |
|  | Total | 22 | 14 | 8 |
| ssDNA viruses | Circoviridae | 1 | 1 | 0 |
|  | Geminiviridae | 0 | 0 | 0 |
|  | Nanoviridae | 0 | 0 | 0 |
|  | Unclassified | 0 | 0 | 0 |
|  | ssDNA viruses | 0 | 0 | 0 |
|  | Total | 1 | 1 | 0 |
| All |  | 83 | 58 | 25 |

\* Two Hepeviridae (ssRNA+).

**Table S2:** Assembly statistics for NCLDV metagenome-assembled genomes and corresponding VIPs

| Metagenome-assembled genome | #contigs | N50 | L50 | Min | Max | Sum | VIPs OTUs<br>(OM-RGC.v1 ID) |
| --- | --- | --- | --- | --- | --- | --- | --- |
| TARA_IOS_NCLDV_Bin_127_6 | 14 | 21,642 | 5 | 8,581 | 35,822 | 267,607 | PolB 000079111 |
| TARA_IOS_NCLDV_Bin_173_3 | 12 | 12,913 | 3 | 2,807 | 34,517 | 108,412 | PolB 000248170 |
| TARA_MED_NCLDV_Bin_284_10 | 34 | 10,936 | 10 | 2,580 | 29,722 | 298,760 | PolB 000328966 |
| TARA_MED_NCLDV_Bin_284_14 | 43 | 14,837 | 11 | 2,756 | 27,607 | 439,843 | PolB 001175669 |
| TARA_IOS_NCLDV_Bin_127_4 | 26 | 5,734 | 10 | 2,560 | 8,505 | 133,765 | PolB 001064263<br>and 010288541 |
| TARA_AON_NCLDV_Bin_289_4 | 17 | 9,468 | 5 | 3,044 | 26,201 | 153,728 | PolB 000200745<br>and 002503270 |
| TARA_MED_NCLDV_Bin_341_10 | 5 | 7,800 | 2 | 2,534 | 7,941 | 30,478 | PolB 002682999 |
| TARA_PON_NCLDV_Bin_65_10 | 35 | 13,866 | 11 | 3,781 | 43,080 | 382,455 | PolB 000079078 |
| TARA_PON_NCLDV_Bin_102_1 | 53 | 4,608 | 18 | 2,606 | 11,485 | 239,832 | PolB 000495602 |
| TARA_AON_NCLDV_Bin_133_8 | 8 | 7,204 | 3 | 2,686 | 10,349 | 51,009 | PolB 000240662 |

N50: The length of the contigs for which half of the assembly size is contained in contigs with a length greater than N50. L50: Number of contigs (or scaffolds) with a size greater or equal to N50.

**Table S3:** Host predictions per viral and host group for 83 VIPs based on phylogeny, co-occurrence analysis, and genomic context

| Virus-host relationship | Positive VIPs | Negative VIPs | Total |
| --- | --- | --- | --- |
| NCLDV-Mamiellales | 10 | 4 | 14 |
| NCLDV-Prymnesiales | 5 | 1 | 6 |
| NCLDV-Pelagophyceae | 2 | 1 | 3 |
| NCLDV-No prediction | 26 | 11 | 37 |
| RNA virus-Copepoda | 7 | 2 | 9 |
| RNA virus-Chaetocerotales | 2 | 0 | 2 |
| RNA virus-Labyrinthulomycetes | 1 | 0 | 1 |
| RNA virus-No prediction | 4 | 6 | 10 |
| ssDNA virus-Copepoda | 1 | 0 | 1 |
| Total | 58 | 25 | 83 |

**Table S4:**
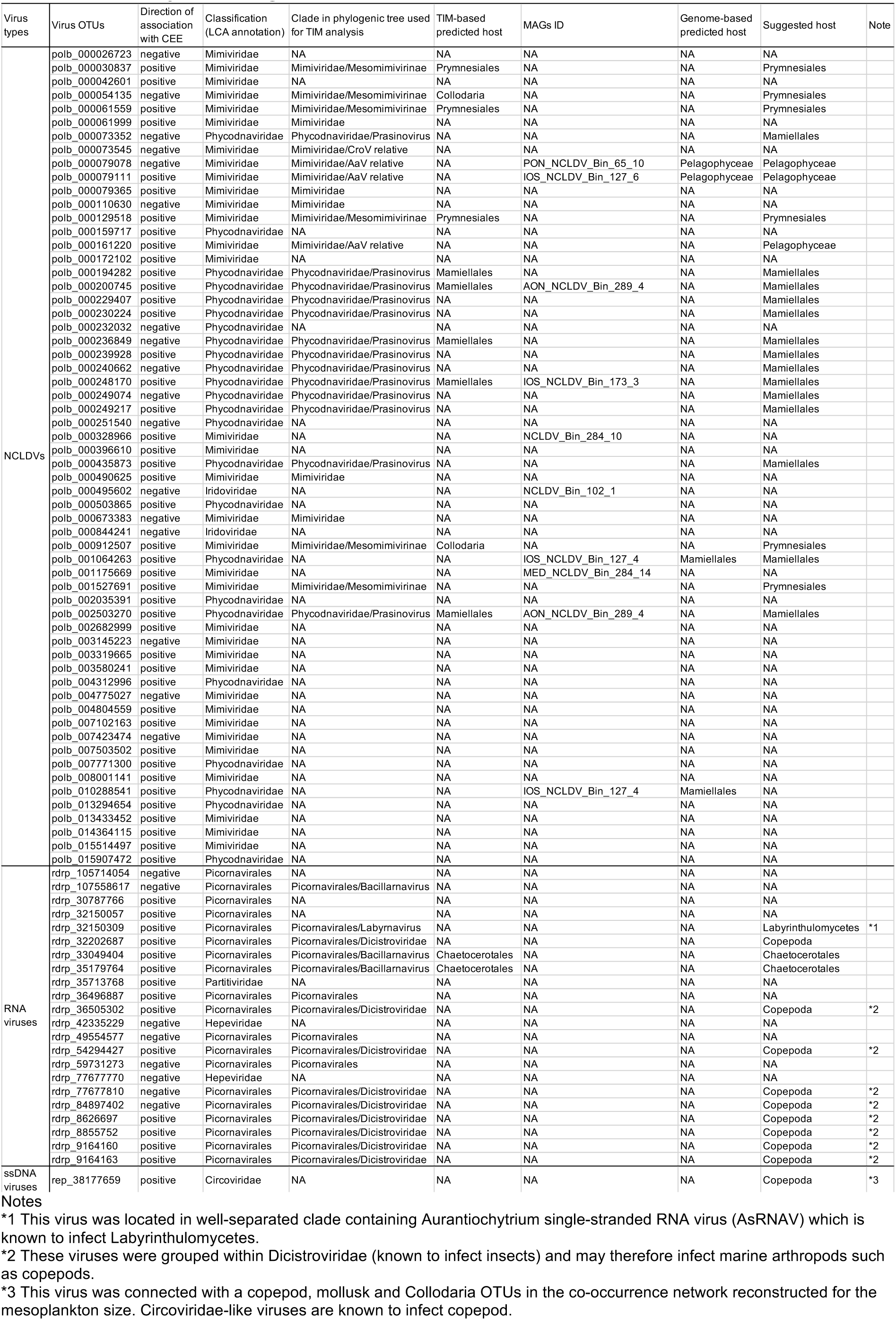
Host prediction per viral OTU for 83 VIPs based on phylogeny, co-occurrence analysis, and genomic context

**Table S5:**
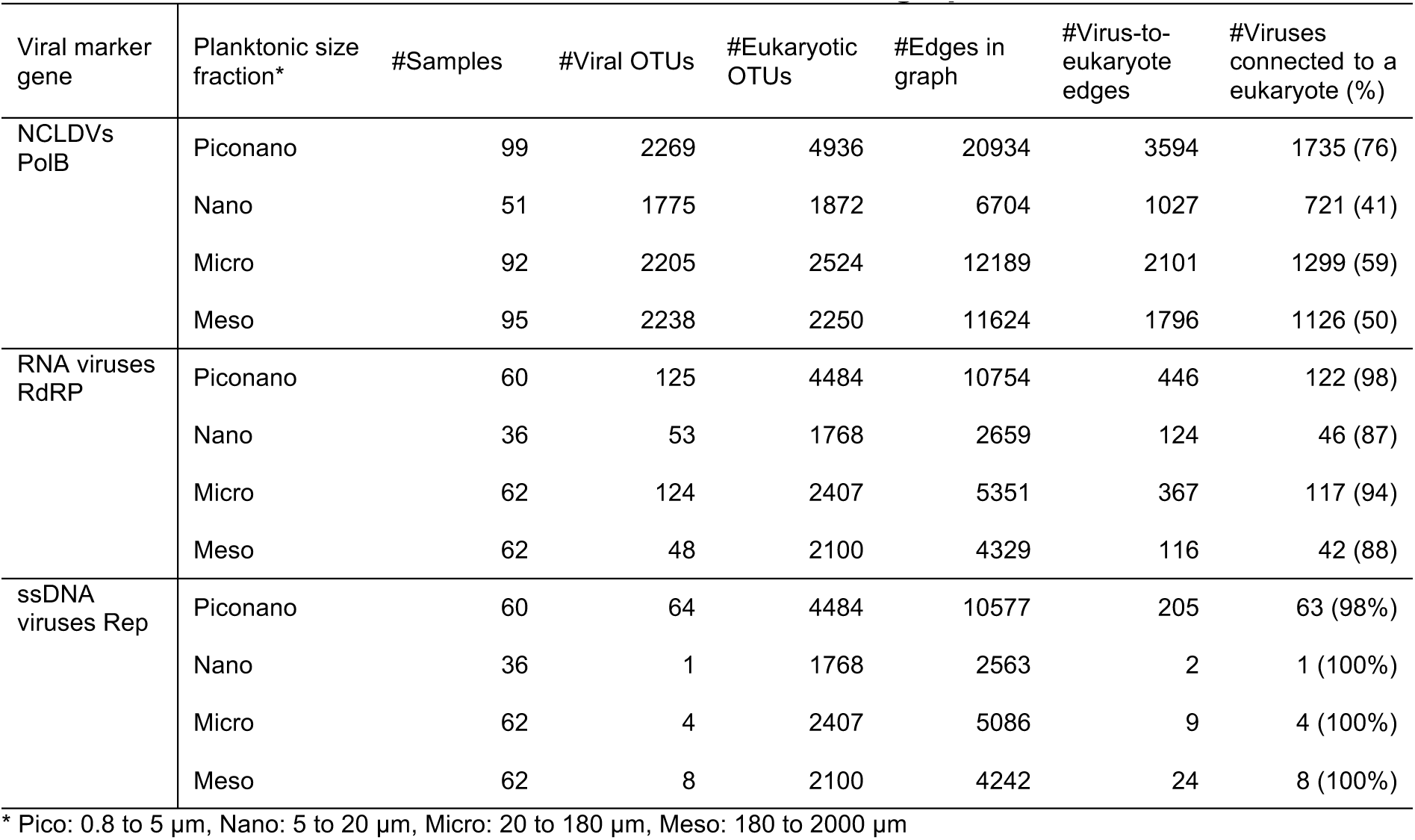
Statistics for the FlashWeave co-occurrence graphs

**Table S6:** Functional differences between eukaryotes found to be best connected to VIPs and non-VIPs

| Functional trait | Positive VIPs (*n* = 50) |  | Non-VIPs (*n* = 983) |  | *P*-value<br>(Fisher's exact<br>test, two<br>sided) | Adjusted *P*-<br>value (BH) (*Q*) |
| --- | --- | --- | --- | --- | --- | --- |
|  | Presence | Absence | Presence | Absence |  |  |
| Chloroplast | 20 | 30 | 276 | 690 | 0.109 | 0.164 |
| Silicification | 11 | 39 | 60 | 920 | 0.000 | 0.001 |
| Calcification | 1 | 49 | 30 | 950 | 1.000 | 1.000 |
| Functional trait | Negative VIPs (*n* = 21) |  | Non-VIPs (*n* = 983) |  | *P*-value<br>(Fisher's exact<br>test, two<br>sided) | Adjusted *P*-<br>value (BH) (*Q*) |
|  | Presence | Absence | Presence | Absence |  |  |
| Chloroplast | 3 | 17 | 276 | 690 | 0.218 | 0.655 |
| Silicification | 0 | 21 | 60 | 920 | 0.632 | 0.947 |
| Calcification | 0 | 21 | 30 | 950 | 1.000 | 1.000 |

**Table S7:** Functional differences between eukaryotes found to be best connected to positive and negative VIPs

| Functional trait | Positive VIPs (n = 50) |  | Negative VIPs (n = 21) |  | *P*-value<br>(Fisher exact<br>test, two<br>sided) | Adjusted *P*-<br>value (BH) (*Q*) |
| --- | --- | --- | --- | --- | --- | --- |
|  | Presence | Absence | Presence | Absence |  |  |
| Chloroplast | 20 | 30 | 3 | 17 | 0.053 | 0.079 |
| Silicification | 11 | 39 | 0 | 21 | 0.027 | 0.079 |
| Calcification | 1 | 49 | 0 | 21 | 1.000 | 1.000 |

## Transparent Methods

### Data context

We used publicly available data generated in the framework of the *Tara* Oceans expedition. Single-copy marker-gene sequences for NCLDVs and RNA viruses were identified from two gene catalogs: the Ocean Microbial Reference Gene Catalog (OM-RGC) and the Marine Atlas of *Tara* Oceans Unigenes (MATOU). The viral marker-gene read count profiles used in our study are as previously reported for prokaryotic-sized metagenomes (size fraction 0.2–3 μm) (Sunagawa et al., 2015) and eukaryotic-sized metatranscriptomes (Carradec et al., 2018). Eukaryotic plankton samples (the same samples were used for metatranscriptomes, metagenomes and 18S rRNA V9-meta-barcodes) were filtered for categorization into the following size classes: piconano (0.8–5 μm), nano (5–20 μm), micro (20–180 μm), and meso (180–2,000 μm). For eukaryotic 18S rRNA V9 OTUs (de Vargas et al., 2015), we used an updated version of the data that included functional trait annotations (chloroplast-bearing, silicified, and calcified organisms) of V9 OTUs. Occurrence profiles are compositional matrices in which gene occurrence are expressed as unnormalized (V9 meta-barcode data) or gene-length normalized (shotgun data) read counts. Indirect measurements of carbon export (mg m^−2^ d^−1^) in 5-m increments from the surface to a 1,000-m depth were taken from Guidi et al. (Guidi et al., 2016) The original measurements were derived from the distribution of particle sizes and abundances collected using an underwater vision profiler. These raw data are available from PANGEA (Picheral et al., 2014). Net primary production (NPP) data were extracted and averaged from 8-day composites of the vertically generalized production model (VGPM) (Behrenfeld and Falkowski, 1997) for the week of sampling. Thus, in this study, the comparisons between NPP and other parameters were not made at the same time point. This might have affected the results of the regression analysis, especially if there were any short-term massive bloom events, although there was no bloom signal during the sampling period.

### Carbon export, carbon export efficiency, and particle size distribution

Carbon flux profiles (mg m^−2^ d^−1^) were estimated based on particle size distributions and abundances. The method used for carbon flux estimation was previously calibrated comparing sediment trap measurement and data from imaging instruments (Guidi et al., 2008). Carbon flux values from depths of 30 to 970 meters were divided into 20-m bins, each obtained by averaging the carbon flux values from the designated 20 m in profiles gathered during biological sampling within a 25-km radius over 24 h when less than 50% of data were missing (Figure S5). Carbon export (CE) was defined as the carbon flux at 150 m (Guidi et al., 2016). Carbon export efficiency was calculated as follows: CEE = CE_deep_/CE_surface_ (Buesseler and Boyd, 2009). To compare stations with different water column structures, we defined CE_surface_ as the maximum CE (in a 20 m window) within the first 150 m. CE_deep_ is the average CE (also in a 20 m window) 200 m below this maximum. The 150 m limit serves as a reference point to automatize the calculation of CE_surface_ and CE_deep_. The 150m-depth layer was selected because often used as a reference depth for drifting sediment trap and because most of the deep chlorophyll maximum (DCM) were shallower except at two (stations 98 (175 m) and 100 (180 m)). The maximum CE_surface_ for these two stations was above 150 m. The sampling strategy of *Tara* Oceans was designed to study a variety of marine ecosystems and to target well-defined meso- to large-scale features (based on remote-sensing data). Therefore, this strategy avoided sampling water with important lateral inputs. Nevertheless, the possibility of having locations with potential lateral transport cannot be excluded.

We obtained the particle size distribution (PSD) profiles generated by the *Tara* Oceans expedition and computed the PSD slope at each depth for all profiles. The slope value (denoted “*b*”) is used as the descriptor of the particle size distribution as defined in a previous work (Guidi et al., 2009). For example, *b* = −5 indicates the presence of a large proportion of smaller particles, whereas *b* = −3 indicates a preponderance of larger particles. We averaged the slope values at each sampling site in the same way as for carbon export flux.

### Identification of viral marker genes from ocean gene catalogs

Viral genes were collected from two gene catalogs: OM-RGC version 1 and MATOU. Sequences in these two gene catalogs are representatives of clusters of environmental sequences (clustered at 95% nucleotide identity). The OM-RGC data were taxonomically re-annotated, with the NCBI reference tree used to determine the last common ancestor modified to reflect the current classification of NCLDVs (Carradec et al., 2018). We automatically classified viral gene sequences as eukaryotic or prokaryotic according to their best BLAST score against viral sequences in the Virus-Host Database (Mihara et al., 2016). DNA polymerase B (PolB), RNA-dependent RNA polymerase (RdRP), and replication-associated protein (Rep) genes were used as markers for NCLDVs, RNA viruses, and ssDNA viruses, respectively. For PolB, reference proteins from the NCLDV orthologous gene cluster NCVOG0038 (Yutin et al., 2009) were aligned using MAFFT-*linsi* (Katoh and Standley, 2013). A hidden Markov model (HMM) profile was constructed from the resulting alignment using *hmmbuild* (Eddy, 2011). This PolB HMM profile was searched against OM-RGC amino acid sequences and translated MATOU sequences annotated as NCLDVs, and sequences longer than 200 amino acids that had hits with *E*-values < 1 × 10^−5^ were selected as putative PolBs. RdRP sequences were chosen from the MATOU catalog as follows: sequences assigned to Pfam profiles PF00680, PF00946, PF00972, PF00978, PF00998, PF02123, PF04196, PF04197, or PF05919 and annotated as RNA viruses were retained as RdRPs. For Rep, we reconstructed an HMM profile using a comprehensive set of reference sequences (Kazlauskas et al., 2018) and searched this profile against the translated MATOU sequences annotated as ssDNA viruses. We kept sequences that had hits with *E*-values < 1 × 10^−5^ and removed those that contained frameshifts. The procedure above identified 3,486 PolB sequences in the metagenomic samples and respectively 975, 388, and 299 RdRP, PolB, and Rep sequences in the metranscriptomes.

### Testing for associations between viruses with CEE, CE_150_, and NPP

To test for associations between occurrence of viral marker genes and CEE, CE_150_, and NPP, we proceeded as follows. Samples with CEE values greater than one and with Z-score greater than two were considered as outliers and removed (this removed the two samples from station 68). Only marker genes represented by at least two reads in five or more samples were retained (lowering this minimal number of required samples down to three or four did not improve the PLS regression model). To cope with the sparsity and composition of the data, gene-length normalized read count matrices were center log-ratio transformed, separately for ssDNA viruses, RNA viruses and NCLDVs. We next selected genes with Spearman correlation coefficients with CEE, CE_150_ or NPP greater than 0.2 or smaller than −0.2 (zero values were removed). To assess the association between these marker genes and CEE, we used partial least square (PLS) regression analysis. The number of components selected for the PLS model was chosen to minimize the root mean square error of prediction (Figure S6). We assessed the strength of the association between carbon export (the response variable) and viral marker genes occurrence (the explanatory variable) by correlating leave-one-out cross-validation predicted values with the measured carbon export values. We tested the significance of the correlation by comparing the original Pearson coefficients between explanatory and response variables with the distribution of Pearson coefficients obtained from PLS models reconstructed based on permutated data (10,000 iterations). We estimated the contribution of each gene (predictor) according to its variable importance in the projection (VIP) score derived from the PLS regression model using all samples. The VIP score of a predictor estimates its contribution in the PLS regression. Predictors with high VIP scores (> 2) were assumed to be important for the PLS prediction of the response variable.

### Phylogenetic analysis

Environmental PolB sequences annotated as NCLDVs were searched against reference NCLDV PolB sequences using BLAST. Environmental sequences with hits to a reference sequence that had > 40% identity and an alignment length greater than 400 amino acids were kept and aligned with reference sequences using MAFFT-*linsi*. Environmental RdRP sequences annotated as were translated into six frames of amino acid sequences, and reference RNA viruses RdRP sequences collected from the Virus-Host Database were searched against the Conserved Domain Database (CDD) using rpsBLAST. The resulting alignment was used to trim reference and environmental RdRP sequences to the conserved part corresponding to the domain, CDD: 279070, before alignment with MAFFT-*linsi*. Rep sequences annotated as ssDNA viruses were treated similarly. PolB, RdRP, and Rep multiple sequence alignments were manually curated to discard poorly aligned sequences. Phylogenetic trees were reconstructed using the the *build* function of ETE3 (Huerta-Cepas et al., 2016) of the GenomeNet TREE tool (https://www.genome.jp/tools-bin/ete). Columns were automatically trimmed using *trimAl* (Capella-Gutiérrez et al., 2009), and trees were constructed using FastTree with default settings (Price et al., 2009).

A similar procedure was applied for the trees used in the hosts prediction analysis albeit selecting sequences for the Phycodnaviridae/Mimiviridae (Figure S8) and the Picornavirales (Figure S9) and removing the ones occurring in fewer than 10 samples, to reduce the size of the tree.

### Virus–eukaryote co-occurrence analysis

We used FlashWeave (Tackmann et al., 2019) with Julia 1.2.0 to predict virus–host interactions based on their co-occurrence patterns. Read count matrices for the 3,486 PolBs, 975 RdRPs, 299 Reps, and 18S rRNA V9 DNA barcodes obtained from samples collected at the same location were fed into FlashWeave. The 18S rRNA V9 data were filtered to retain OTUs with an informative taxonomic annotation. The 18S rRNAV9 OTUs and viral marker sequences were further filtered to conserve only those present in at least five samples. FlashWeave networks were learned for each of the four eukaryotic size fractions with the parameters ‘heterogenous’ = false and ‘sensitive’ = true, and edges receiving a weight > 0.2 and a *Q*-value < 0.01 (the default) were retained. The number of samples per size fraction ranged between 51 and 99 for NCLDVs and between 36 and 62 for RNA and ssDNA viruses. The number of retained OTUs per size fraction varied between 1,775 and 2,269 for NCLDVs and between 48 and 125 for RNA viruses (Table S5).

### Mapping of putative hosts onto viral phylogenies

In our association networks, individual viral sequences were often associated with multiple 18S rRNA V9 OTUs belonging to drastically different eukaryotic groups, a situation that can reflect interactions among multiple organisms but also noise associated with this type of analysis (Coenen and Weitz, 2018). To extract meaningful information from these networks, we reasoned as follows. We assumed that evolutionarily related viruses infect evolutionarily related organisms, similar to the case of phycodnaviruses (Clasen and Suttle, 2009). In the interaction networks, the number of connections between viruses in a given clade and the associated eukaryotic host group should accordingly be enriched compared with the number of connections with non-host organisms arising by chance. Following this reasoning, we assigned the most likely eukaryotic host group as follows. The tree constructed from viral marker-gene sequences (PolB, RdRP or Rep) was traversed from root to tips to visit every node. We counted how many connections existed between leaves of each node and the V9-OTUs of a given eukaryotic group (order level). We then tested whether the node was enriched compared with the rest of the tree using Fischer’s exact test and applied the Benjamini–Hochberg procedure to control the false discovery rate among comparisons of each eukaryotic taxon (order level). To avoid the appearance of significant associations driven by a few highly connected leaves, we required half of the leaves within a node to be connected to a given eukaryotic group. Significant enrichment of connections between a virus clade and a eukaryotic order was considered to be indicative of a possible virus–host relationship. We refer to the above approach, in which taxon interactions are mapped onto a phylogenetic tree of a target group using the organism’s associations predicted from a species co-occurrence-based network, as TIM, for Taxon Interaction Mapper. This tool is available at https://github.com/RomainBlancMathieu/TIM. This approach can be extended to interactions other than virus–host relationships.

### Assembly of NCLDV metagenome-assembled genomes (MAGs)

NCLDV metagenome-assembled genomes (MAGs) were assembled from *Tara* Oceans metagenomes corresponding to size fractions > 0.8 μm. Metagenomes were first organized into 11 ‘metagenomic sets’ based upon their geographic coordinates, and each set was co-assembled using MEGAHIT (Li et al., 2015) v.1.1.1. For each set, scaffolds longer than 2.5 kbp were processed within the bioinformatics platform anvi’o (Eren et al., 2015) v.6.1 following methodology described previously for genome-resolved metagenomics (Delmont et al., 2018). Briefly, we used the automatic binning algorithm CONCOCT (Alneberg et al., 2014) to identify large clusters of contigs using both sequence composition and differential coverage across metagenomes within the set. We then used HMMER (Eddy, 2011) v3.1b2 to search for a collection of eight NCLDV gene markers (Guglielmini et al., 2019), and identified NCLDV MAGs by manually binning CONCOCT clusters of interest using the anvi’o interactive interface. The interface displayed hits for the eight gene markers alongside coverage values across metagenomes and GC-content. Finally, NCLDV MAGs were manually curated using the same interface, to minimize contamination as described previously (Delmont and Eren, 2016).

### Taxonomic composition of genes predicted in NCLDV genomes of VIPs

VIP’s PolB sequences were searched (using BLAST) against MAGs reconstructed from the metagenomes of the eukaryotic size fraction (> 0.8 μm) and against contigs used to produce OM-RGCv1. Genome fragments covering 95% of the length of PolB VIPs with > 95% nucleotide identity were considered as originating from a same viral OTUs. Genes were predicted and annotated taxonomically with the same procedure described above (identification of viral marker genes). Genes contained in viral genome fragments and annotated as cellular organisms with amino acid identities > 60% were manually inspected (Supplemental Data 2).

### Statistical test

All the statistical significance assessments were performed with two-sided test.

## Notes

ftp://ftp.genome.jp/pub/db/community/tara/Cpump/

### Competing Interest Statement

The authors have declared no competing interest.

### Summary of Updates

The definition of CEE has been changed. NCLDV MAG analysis has been added.

